# Cep55 regulation of PI3K/Akt signaling is required for neocortical development and ciliogenesis

**DOI:** 10.1101/2021.01.08.425857

**Authors:** Behnam Rashidieh, Belal Shohayeb, Amanda Louise Bain, Patrick R. J. Fortuna, Debottam Sinha, Andrew Burgess, Richard Mills, Rachael C. Adams, J. Alejandro Lopez, Peter Blumbergs, John Finnie, Murugan Kalimutho, Michael Piper, James Edward Hudson, Dominic Ng, Kum Kum Khanna

## Abstract

Homozygous nonsense mutations in CEP55 are associated with several congenital malformations that lead to perinatal lethality suggesting that it plays a critical role in regulation of embryonic development. CEP55 has previously been studied as a critical regulator of cytokinesis predominantly in transformed cells and its deregulation is linked to carcinogenesis. However, its molecular functions during embryonic development in mammals have not been clearly defined. We have generated a Cep55 knockout (Cep55^-/-^) mouse model which demonstrated perinatal lethality associated with a wide range of neural defects. Focusing our analysis on the neocortex, we show that Cep55-/- embryos exhibited depleted neural stem/progenitor cells in the ventricular zone as a result of significantly increased cellular apoptosis. Mechanistically, we demonstrated that Cep55-loss downregulates the pGsk3β/β-Catenin/Myc axis in an Akt-dependent manner. The phenotype was recapitulated using human cerebral organoids and we could rescue the phenotype by inhibiting active Gsk3β. Additionally, we show that Cep55-loss leads to a significant reduction of ciliated cells, highlighting its novel role in regulating ciliogenesis. Collectively, our findings demonstrate a critical role of Cep55 during brain development and provide mechanistic insights that may have important implications for genetic syndromes associated with Cep55-loss.

## Introduction

Centrosomal protein 55 kDa (CEP55) is a crucial regulator of cytokinesis, the final stage of mitotic cell division[1]. CEP55 is highly upregulated in a wide spectrum of tumors and has been reported to play critical roles in the regulation of the *PI3K/AKT* pathway, stemness, genomic stability, and cell cycle progression [2,3]. Despite extensive investigation on the roles of human CEP55 in tumorigenesis, its physiological role during development has remained largely uncharacterized. Recently, germline mutations of CEP55 in humans have been described in two lethal *CEP55*-associated syndromes, Meckel-Gruber syndrome (MKS)-like Syndrome[4,5] and MARCH (Multinucleated neurons, Anhydramnios, Renal dysplasia, cerebral hypoplasia, and Hydranencephaly) [6]. These syndromes exhibit multiple severe clinical manifestations including several congenital malformations that lead to perinatal lethality. Homozygous nonsense mutations in *CEP55* that are predicted to lead to loss of protein were identified in affected fetuses. However, mechanisms underlying complex Cep55-deficient developmental phenotypes remain elusive. We have generated a Cep55 knockout (KO) mouse model to reveal developmental phenotype and to explore whether it recapitulates the clinical condition. Additionally, we have used cerebral organoids generated from pluripotent stem cells as a promising approach to investigate the mechanism of Cep55-associated neurodevelopment phenotype.

By generating a mouse model and human cerebral organoids lacking Cep55, here we found that *Cep55* deletion resulted in a reduction in the size of mouse brains and human cerebral organoids due to excessive apoptosis of neural progenitor cells (NPC). Additionally, we discovered a critical role for Cep55 in regulating cilia formation. Mechanistically, we show for the first time that Cep55 regulates neural development through the Akt-downstream effector, Gsk3β, and its mediators β-Catenin and Myc which are known regulators of neural proliferation and differentiation [7]. However, Cep55 regulation of ciliogenesis occurs through AKT independent of Gsk3β. Together, these results illustrate an important role of Cep55 in regulating neurogenesis and ciliogenesis in an Akt dependent manner in mice.

## Results

### Loss of Cep55 leads to perinatal lethality in mice

To investigate the physiological role of CEP55 during development, we generated a KO mouse model of *Cep55* using the “KO first” allele design wherein the targeted allele acts as a gene-trap to form a non-functional allele (fig. S1A). Correct targeting was validated independently by genotyping PCR alongside *Cep55* transcript and protein expression using RT-qPCR and immunoblotting analysis, respectively (fig. S1B-D). To generate the colony of *Cep55^-/-^* (KO) mice, we intercrossed *Cep55^+/-^* mice, with the expectation that approximately 25% of the offspring would be of a *Cep55^-/-^* genotype according to Mendelian ratios. Interestingly, after genotyping more than 77 offspring from these breedings across 19 litters, we could not detect any viable *Cep55^-/-^* mice, indicating that genetic loss of *Cep55* led to embryonic or perinatal lethality (Sup table 1). To define the time point of lethality, pregnant dams from *Cep55^+/-^* intercrosses were euthanized at different stages of pregnancy, ranging from E11.5 - E18.5, and embryos collected for phenotypic evaluation. Interestingly, we were able to obtain viable *Cep55^-/-^* offspring at each gestational stage (E11.5-E18.5) except at the time of birth (Sup table 2). Notably, embryos collected at both E14.5 and E18.5 exhibited significant dwarfism, based on crown-rump length measurements, when compared to control (*Cep55^+/+^*) embryos. Additionally, *Cep55^-/-^* embryos exhibited an increased thickness of the neck and a flattened head (Fig. 1A, B).

**Figure 1.**
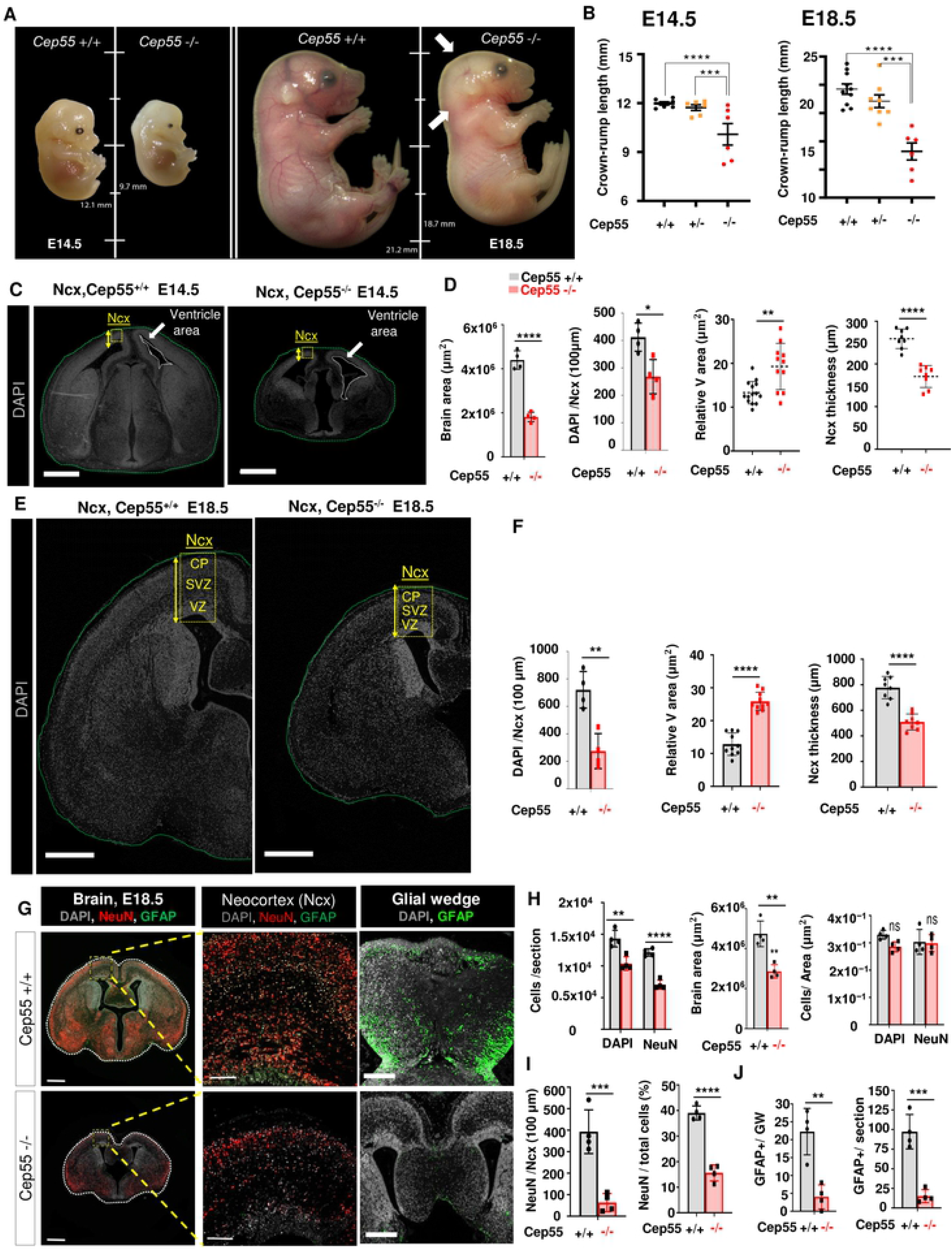
Loss of *Cep55* leads to perinatal lethality and microcephaly in mice. **(A)** Comparison of size (length, mm) and morphology of E14.5 (left) and E18.5 (right) *Cep55*^+/+^ and *Cep55*^-/-^ embryos. **(B)** Comparison of the crown-rump length (mm) of E14.5 (left) and E18.5 (right) *Cep55*^+/+^ and *Cep55*^-/-^ embryos. Data represent the mean ± SD, n = 6–10 embryos per genotype. **(C,E)** Representative images of *Cep55*^+/+^ (left) and *Cep55^-/-^* (right) neocortices (Ncx, yellow box) showing relative size of the the Ncx. The yellow two sided arrow represents the thickness of the Ncx, green dashed line shows the brain area, white dashed line shows the ventricle area and the yellow box resembles the analysis area, **(C)** Representative images of *Cep55^+/+^* and *Cep55^-/-^* mouse brains at E14.5, scale= 50μm. **(D)** Comparison of brain area of *Cep55^+/+^* and *Cep55^-/-^* E14.5 embryonic brains (left), and quantification of brain cell density (DAPI count within 100μm^2^ area of the neocortex) (Middle left) Quantification of the relative ventricle area (μm^2^; total area shown/total brain area) (Middle right), and Ncx thickness (right). **(E)** Representative images of *Cep55*^+/+^ and *Cep55^-/-^* mouse brains at E18.5, scale= 700μm. **(F)** Comparison of *Cep55*^+/+^ and *Cep55^-/-^* for left: total DAPI+ cell count in the Ncx; middle: relative ventricle area (μm2; total ventricular space/total brain area) and right: Ncx thickness. **(G)** Representative images of whole coronal section (left), boxed region at increased magnification (middle), and medial region/glial wedge (right) of E18.5 *Cep55^+/+^* (upper) and *Cep55^-/-^* (lower) embryonic mouse brains. Images show staining for NeuN (neuronal nuclei, mature neurons, red) and GFAP (Glial fibrillary acidic protein, marks astrocytes and ependymal cells, green). Right panel (glial wedge) shows the population of mature radial glia. Corpus callosum dysgenesis in Cep55^-/-^ brain (lower right), the boxed area in this panel shows GFAP expression in a magnified zone of glial wedge. Scale= 600 μm (left), Scale= 100 μm (middle), Scale= 400μm (right). **(H)** Comparison of *Cep55*^+/+^ and *Cep55^-/-^* embryonic brain overall cell and neuron number per section (left), brain area (middle), and cell/neuron density (right). N=4, P<0.0036, 0.0001. **(I)** Comparison of NeuN-positive neurons normalized to 100μm neocortical area (Ncx) in *Cep55^+/+^* and *Cep55^-/-^* E18.5 embryonic brains (left) and percentage of NeuN-positive cells across whole brain section normalized to the total number of cells identified by DAPI fluorescence (right). Data represent mean + SD across two regions from n=4 independent embryos per genotype. **(J)** Quantification of GFAP-positive cells in the glial wedge of *Cep55^+/+^* and *Cep55^-/-^* E18.5 embryonic brains (left) and GFAP-positive cells in the whole section (right). Data represent mean ± SD of four embryos, N=4, average count of duplicate technical repeats, Student’s t-test, *P < 0.05, **P < 0.01, ***P < 0.001, ****P < 0.0001).

To determine if the loss of a single allele of *Cep55* would cause phenotypic changes, we performed the histological examination of multiple organs from eight-week-old *Cep55^+/-^* mice relative to *Cep55^+/+^* littermates. We observed no significant differences in the pathohistology or size of respective organs (fig S1E) indicating that loss of a single allele of *Cep55* does not impact physiological development. Additionally, the monitoring of both genotypes showed no significant differences in body weight for the first 20 weeks (fig S1F). These data suggest that a single allele of *Cep55* is largely sufficient to maintain physiological functions.

Several recent reports have shown that CEP55 functional loss in humans leads to a range of congenital abnormalities, all with defective brain development[4–6]. Therefore, we next sought to examine the expression pattern of *Cep55* using single-cell transcriptomic data of mouse neocortical development[8]. This analysis revealed that *Cep55* expression levels are highest in the NP cells of E14 embryos (fig S1G). Moreover, investigating human fetal brain data based on Allen brain atlas revealed that expression of *Cep55* peaks from weeks 8-10 of gestation, followed by a reduction after 16 weeks, minimal detection between weeks 27-35, and becoming detectable again three weeks prior to birth[9]. This expression pattern corresponds with the timing of human neurogenesis in the neocortex through neurogenic divisions and neuronal differentiation from radial glial cells (RGCs)[10]. To validate the expression of *Cep55* during development, we performed β-galactosidase staining in the *Cep55^+/-^* mice, where the targeted allele contains a LacZ reporter. In the isolated brain of mouse embryos, a gradient of expression of CEP55 in the neocortex was detected at E12.5, diminishing at E14.5 to become undetectable at E16.5 (fig S1H). Collectively, our data suggest that *Cep55* plays a critical role during embryogenesis particularly neurogenesis but is dispensable for adult tissue homeostasis.

### *Cep55* loss causes gross morphological defects in mouse embryos

Given the neurodevelopmental expression pattern of Cep55, we predicted significant neural deficits would arise from Cep55 loss. To investigate this, we first performed hematoxylin and eosin (H&E) staining of sagittal and coronal sections of E18.5 embryos of *Cep55^+/+^, Cep55^+/-^* and *Cep55^-/-^* mice. We observed no gross morphological differences in the lung, intestine or liver among the respective genotypes. However, we noted prominent abnormalities in the brain of *Cep55^-/-^* embryos when compared to respective controls, which were characterized by a partial failure (hypoplasia) and disorder (dysplasia) of normal structural brain development. The cerebellum was hypoplastic, with marked thinning of the germinative external-granular layer (EGL) and a diminution and disorganization of neurons (fig S2A, Right). In addition, the neuronal population of the olfactory bulb was disorganized and depleted (fig S2A,,Left). Furthermore, coronal sectioning of the brain revealed neocortical depletion of neurons and ventricular dilatation, as well as smaller germinal regions in both dorsal and ventral telencephalon (fig. S2B). The neocortices of *Cep55^-/-^* brains were hypoplastic and dysplastic, with diminished and disorganized neurons. In addition to apoptosis in the neocortex, there were also multifocal areas of cortical necrosis and parenchymal loss, with evidence of phagocytosis of affected neurons by macrophage-like cells (fig. S2C, upper). Numerous bi-nucleated neurons were also found in the neocortex of *Cep55*^-/-^ (fig. S2C, lower). To measure this defect, we stained mature neurons with NeuN (RBFOX3) and quantified the number of bi-nucleated neurons in the neocortex of brain.The proportion of multinucleated neurons immunostained by NeuN in the cortical region of *Cep55^-/-^* mice was increased compared to that of *Cep55*^+/+^ (fig. S2D,E), a phenotype reminiscent of the described changes in human embryos with MARCH syndrome[6].

As neurogenesis in neocortical layers peaks at approximately E14.5[11,12] and the highest expression of *Cep55* was found at this embryonic stage (fig S1H), we chose this gestational stage for characterizing the phenotype and cellular behavior. Also, to better characterize the specific disruption to neural cells, we focused our investigation on the neocortex, a well-characterized region of the developing forebrain with prominent CEP55 expression. Strikingly, brain sizes of *Cep55^-/-^* E14.5 embryos were found to be significantly smaller compared to that of *Cep55^+/+^* by measuring the brain area (Fig. 1C,D). We also found a reduced number of cells in the neocortex of *Cep55^-/-^* embryos. Furthermore, the size of the ventricle relative to the total brain area was larger and dilated in *Cep55^-/-^* mice, consistent with our previous histopathology observations. We also observed that the thickness of the cortex was reduced in *Cep55^-/-^* brains compared to *Cep55*^+/+^ (Fig. 1C,D). Similar to our observations at E14.5, we noted fewer cells (DAPI) in the cortex of *Cep55^-/-^* embryos when compared to *Cep55*^+/+^ at E18.5 (Fig. 1E,F). Consistently, the ventricles were larger and dilated in *Cep55^-/-^* brains and cortex thickness was reduced (Fig. 1E,F). Also, brain sizes of *Cep55^-^* E18.5 embryos were found to be significantly smaller compared to that of *Cep55^+/+^* (Fig. 1G,H). Interestingly, this size reduction (*Cep55*^-/-^ brain area) likely resulted from decreases in the number of both total cells (DAPI stained) and neurons (NeuN stained), since the density of the cells (cells/area) after normalization to total brain area was not significantly different between genotypes (Fig. 1G,H).

To further investigate the reduction of neurons, we stained brain sections for markers of distinct populations including NeuN for mature neurons and glial fibrillary acidic protein (GFAP) to mark astrocytes. The mature neurons (NeuN-positive cells) were reduced in numbers in the neocortex in *Cep55^-/-^* compared to that of *Cep55^+/+^* embryos, even after normalization to the total cell number as assessed by DAPI staining (Fig. 1G,I). Interestingly, GFAP-positive cells in the neocortex were reduced in *Cep55^-/-^* embryos when compared to *Cep55^+/+^* (Fig. 1G, J), suggesting potential defects in the central nervous system development. In the cortical region, GFAP can be a marker of either astrocytes or the radial-glial-like neuronal stem cells and represents mature radial glia which can be seen in the medial region (e.g. glial wedge). We also observed a drastic reduction in GFAP-expressing cells at the cortical midline, neocortex, and whole section in *Cep55^-/-^* brain compared to that of *Cep55^+/+^*. These cells are critical in facilitating the crossing of axons through the corpus callosum[13]. In line with the lack of GFAP-expressing cells, at the midline, we observed dysgenesis of the corpus callosum in mutant mice at this age (Fig. 1G). Taken together, these data show that loss of *Cep55* results in defective neuropathological phenotypes in mice.

### Cep55 regulates the fate of radial glial and intermediate progenitor cells

As we found a reduced number of both neurons and astrocytes, we next sought to determine how *Cep55* regulates NP differentiation and development during neurogenesis, with a focus on the different neuroepithelial layers of the neocortex during embryonic development. Neurogenesis in the developing neocortex occurs with the contribution of two types of NPs: radial glial cells (RGCs) and intermediate progenitor cells (IPCs)[14]. The former produce neurons and glia which divide at the ventricular zone (VZ; the apical surface), and express the homeodomain transcription factor, PAX6. The latter, which are derived from radial glial cells, produce only neurons, divide within the basally located subventricular zone (SVZ) and express TBR2, a T-domain transcription factor. The subsequent transition from IPCs to postmitotic projection neurons (PMN) in the cortical plate (CP) is marked by the onset of TBR1 expression[14]. We next, investigated the different populations of progenitor cells within the nascent cortex, to determine how a deficit in cortical neuron number might arise. We categorized RGCs as the PAX6^+^ TBR2^-^ population, as some newborn TBR2^+^ IPCs retain PAX6 expression transiently[15]. Immunostaining for PAX6, TBR2, and TBR1 at E14.5 revealed a reduction in the number of RGCs, IPCs, and post-mitotic neurons as a proportion of the total cortical cell number in *Cep55^-/-^* mice compared to Cep55^+/+^ (Fig. 2A, B). Accordingly, we also observed a reduced population of neurons marked by Tuj-1, the neuron-specific class III β-tubulin in *Cep55^-/-^* neocortices compare to *Cep55^+/+^* (Fig 2C). Next, we investigated how *Cep55* loss impacts mitosis, proliferation, and apoptosis of NPCs by staining for phosphohistone H3-S10 (pH3), Ki67, and TUNEL, as markers of these cellular processes, respectively. We found reduced proportion of pH3 positive-cells in *Cep55^-/-^* mice, when pH3-positive cell pool was normalized to the total number of cells, indicative of either mitotic defects or delayed mitosis/cytokinesis in the absence of *Cep55* (Fig 2D). Consistently, the proliferation index at E14.5 was significantly reduced in *Cep55^-^* animals (Fig. 2E). Finally, we found high levels of apoptosis as marked by TUNEL staining in *Cep55*^-/-^ neocortices compared to controls at E14.5 when assessed as a percentage of total cell number (Fig 2F). Interestingly, Western blot analysis of mouse embryonic brain (E14.5) extract also showed upregulation of cleaved caspase-3 in *Cep55^-/-^* brains consistent with IHC results and histopathological observations (fig S2F). Collectively, these data suggest that reductions in NPCs are due to elevated levels of cell death and reduced proliferation. Our data indicate a role for *Cep55* in the survival and viability of neural progenitor populations in neocortical development.

**Fig 2.**
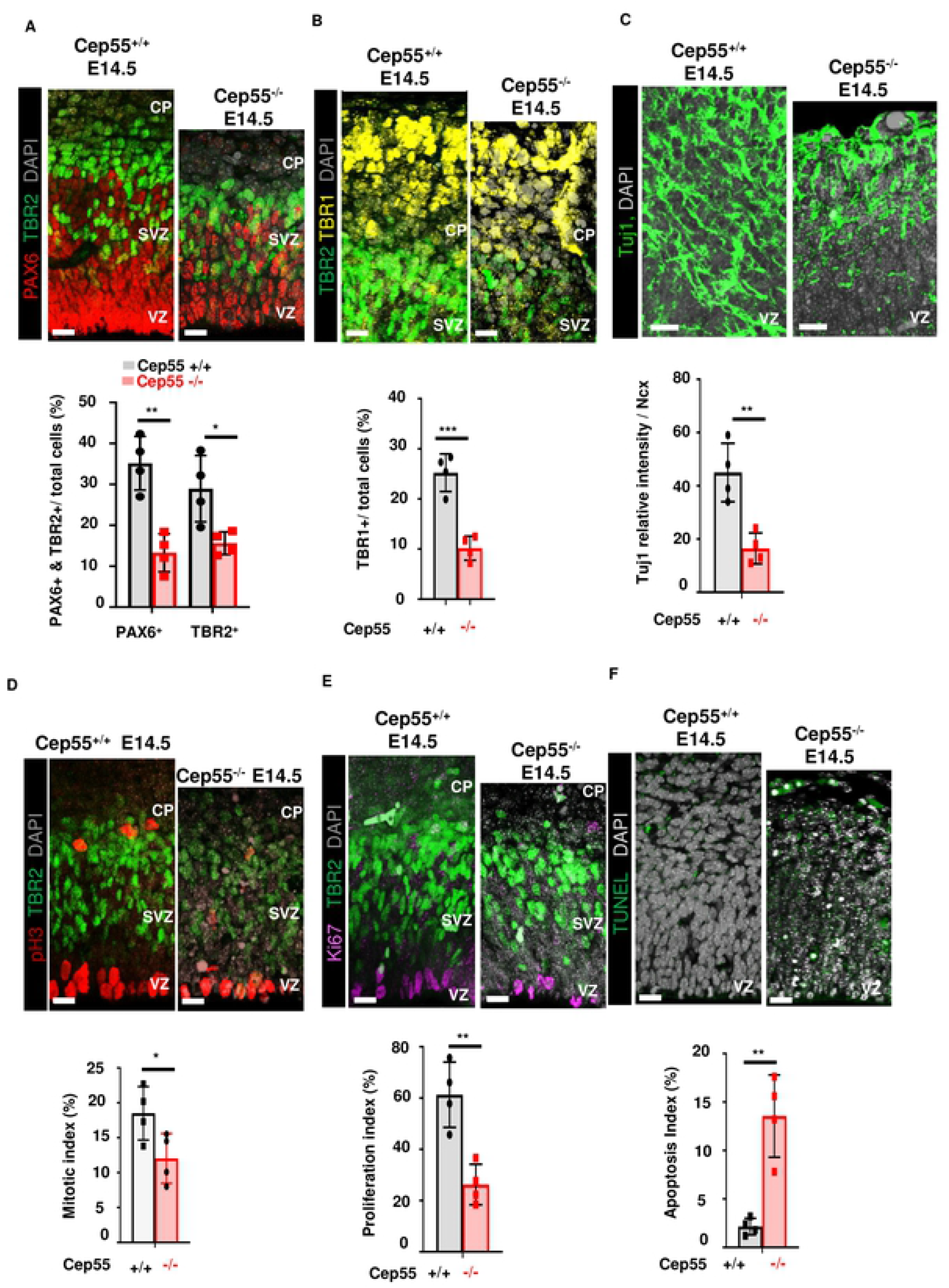
*Cep55* regulates cell fate of radial glial and intermediate progenitor cells, and neurons. **(A)** Representative images of radial glial cells (RGC; PAX6+) at VZ, intermediate progenitor cells (IPC; TBR2+) at SVZ and total cells (DAPI) at E14.5, scale = 15 μm (upper panel). Quantification of percentage of RGCs (PAX6+) and IPCs (TBR2+) in *Cep55^+/+^* and *Cep55^-/-^* neocortices (Ncx) at E14.5 (lower). **(B)** Representative images of IPCs (TBR2+) at SVZ, post-mitotic neurons (TBR1+) at CP and total cells (DAPI, gray) at E14.5, scale = 15 μm (upper), quantification of the percentage of post-mitotic-neurons (TBR1+) in *Cep55*^+/+^ and *Cep55^-/-^* neocortices at E14.5 (lower). **(C)** Neuron-specific class III β-tubulin (Tuj1), relative intensity of Tuj1 staining quantified within a 100μm^2^ field of view. **(D)** Phosphohistone H3 (pH3; mitotic cells) in the VZ and SVZ, co-stained with TBR2 to identify proliferating IPCs at E14.5, scale = 15 μm. Quantification of the percentage of total cells expressing pH3 to show the mitotic index. **(E)** Proliferating cells (Ki67+), IPCs (TBR2+, green) delineating the SVZ, and total cells (DAPI) at E14.5, scale = 15 μm (upper), Comparison of the proportion of Ki67+ cells in *Cep55*^+/+^ and *Cep55^-/-^* neocortices to show proliferation index (lower). **(F)** Apoptotic cells (TUNEL) and total cells (DAPI) in the Ncx, at E14.5, scale = 15 μm (upper), comparison of proportion of apoptotic cells in *Cep55^+/+^* and *Cep55^-/-^* neocortices to show apoptosis index (lower).

### *Cep55* knockdown induces apoptosis in radial glial cells of human cortical organoid

We next investigated the effect of *CEP55*-loss in human cerebral brain organoids generated from embryonic pluripotent stem cells (HES3). Cerebral organoids mimic the unique and dynamic features of early human cortical development in culture, enabling detailed analysis of organ pathogenesis due to particular genetic deregulation or dysfunction[16].

We performed knock-down of Cep55 in differentiated HES3 organoid cultures using adenoviral GFP-tagged shRNA against CEP55 or scrambled control (U6). Knockdown was performed after cerebral organoid induction to circumvent potential apoptosis caused by *CEP55* loss. Within 24-48 hours of adenoviral transduction, we observed a significantly increased level of the integrated virus as marked by GFP expression for control CMV-eGFP and sh-CEP55 (shRNA U6 scrambled control did not contain a GFP tag) (Fig. 3A). The knock-down of CEP55 (Fig. 3B) led to a decrease in the overall size of the transfected organoid, which is in line with the observed microcephaly phenotype (data not shown). Given the extent of apoptosis observed in the mouse model, we characterized the effect of CEP55 knock-down in the cerebral organoids after 24 hours of infection. A significant reduction of PAX6-positive NPs was observed within 24 hours of *CEP55* shRNA transduction in organoids compared to control (Fig. 3C, D). However, no significant difference was found in pH3 positive cells (Fig. 3C, E). The reduced number of PAX6 cells was likely caused by a dramatic increase in cell death (cleaved caspase 3) in organoids transduced with *CEP55* shRNA (Fig. 3F, G). Consequently, we observed a significant decrease in TUJ1 (class III beta-tubulin) positive neurons between controls and *CEP55*-knockdown organoids (Fig. 3F, H). Collectively, our findings of reduced RGC numbers from apoptotic cell death in human cerebral organoids are consistent with the *Cep55^-/-^* phenotype observed in mice.

**Fig 3.**
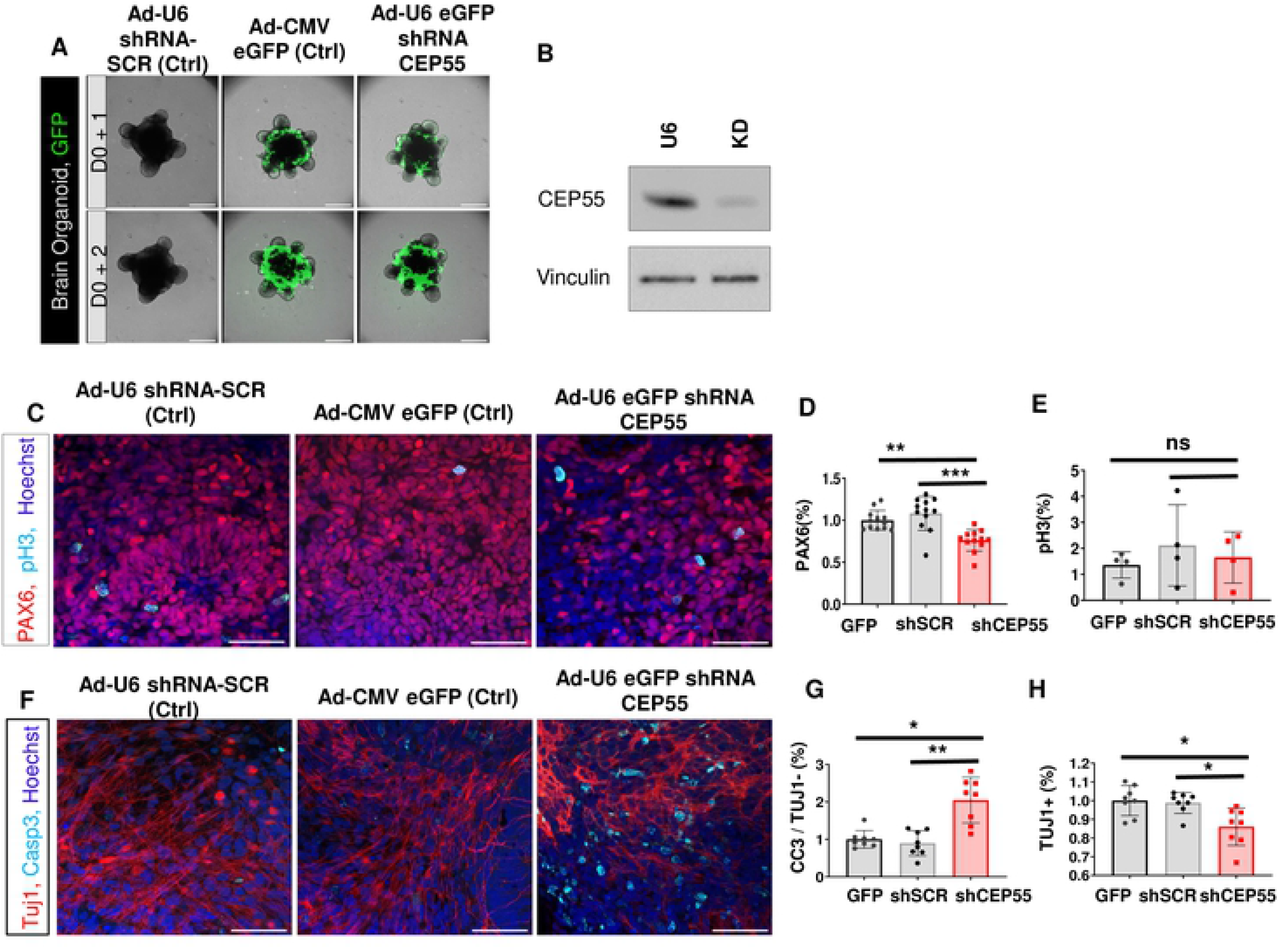
*CEP55* knockdown induces cell death in neural progenitors of human cerebral organoid. **(A)** Day 16 of human iPSC-derived cerebral brain organoids infected with U6 scrambled control (left), CMV GFP control (middle) and CEP55 knockdown (KD) (right) adenoviral shRNAs with GFP-tag. Expression levels of integrated viral GFP increased significantly between days 1 to 2 post-infection at a MOI of 10. shRNA U6 scrambled control did not contain a GFP tag. Upper panel: D0 (infection day) +1; lower panel: D0 +2, scale bars= 50μm, immunofluorescent labeling of cerebral organoids was done after 24 hours of shRNA infection. **(B)** Immunoblot shows knockdown of CEP55 in adenoviral shRNA against CEP55 transduced organoid compare to U6 scrambled control. Vinculin = loading control. **(C)** Representative images of labeling for neural progenitor PAX6, mitotic marker pH3 and nuclear Hoechst showed a decrease in neural progenitor PAX6 in the CEP55 KD shRNA infected organoids compared to the U6 and GFP controls. **(D-E)** Quantified immunolabelled organoids (normalized to CMV GFP control) for **(D)** PAX6 and **(E)** pHH3. **(F)** Representative images of labeling for neural-specific tubulin TUJ1, apoptotic marker cleaved caspase 3, and nuclear Hoechst showed a clear increase in cell death in the CEP55 KD samples compared to the controls. **(G-H)** Quantified immunolabelled organoids (normalized to CMV GFP control) for **(G)** Cleaved Caspase-3 in non Tuj1+ cells **(H)** Total Tuj1+ cells. For all calculations, data represented mean ± SD number of organoids indicated in figures. Non-parametric one-way ANOVA performed; ** p < 0.01, scale = 50 μm.

### Cep55^-/-^ mice exhibit cilial abnormalities

The findings presented above using mouse embryos and human cerebral organoids clearly document the role of *Cep55* in neural development. Notably, nonsense truncating mutations in CEP55 are associated with MKS-like syndrome, a lethal fetal ciliopathy[4,5]. Primary cilia (cilia, hereafter) perform important functions in neurodevelopment, are localized to and extend from RGCs into the lateral ventricle, and are also present in other NPCs and neuron populations[17]. Dysfunction of the ciliary axoneme, basal body, or cilia anchoring structures can all cause defects in cilia organization, leading to ciliopathies[18]. Next, we investigated the involvement of Cep55 in the regulation of ciliogenesis in the developing neocortex at E14.5 (fig S3A) and E18.5 (Fig 4A,B). We performed immunostaining on the embryonic mouse brain sections from *Cep55^+/+^* and *Cep55^-/-^* mice using Arl13b (a marker of ciliary membranes), γ-tubulin (basal body), DAPI (DNA marker), and *Cep55* to evaluate and compare their expression and localization. We observed fewer cilia in the VZ of *Cep55^-/-^* brains at E14.5 and E18.5 compared to *Cep55^+/+^* brain. Our analysis revealed a decrease in both number and percentage of ciliated cells throughout the cortical layers at both E14.5 and E18.5, particularly in apical progenitors localized in the ventricle membrane of *Cep55^-/-^* compared to *Cep55^+/+^* brains (Fig 4A,B and fig S3A). This decrease in ciliated cells in the SVZ was independent of the reduction of IPCs, as we counted ciliated TBR2^+^ cells (fig S3B). Consistently, the ratio of ciliated RGCs (PAX6^+^) cells decreased significantly in *Cep55^-/-^* compared to control which is indicative of cilia defects independent of RGC population drop (fig S3B). Given the important role of cilia during neurodevelopment, we further examined a potential role of *Cep55* in regulation of ciliogenesis in an *in vitro* model.

**Fig 4.**
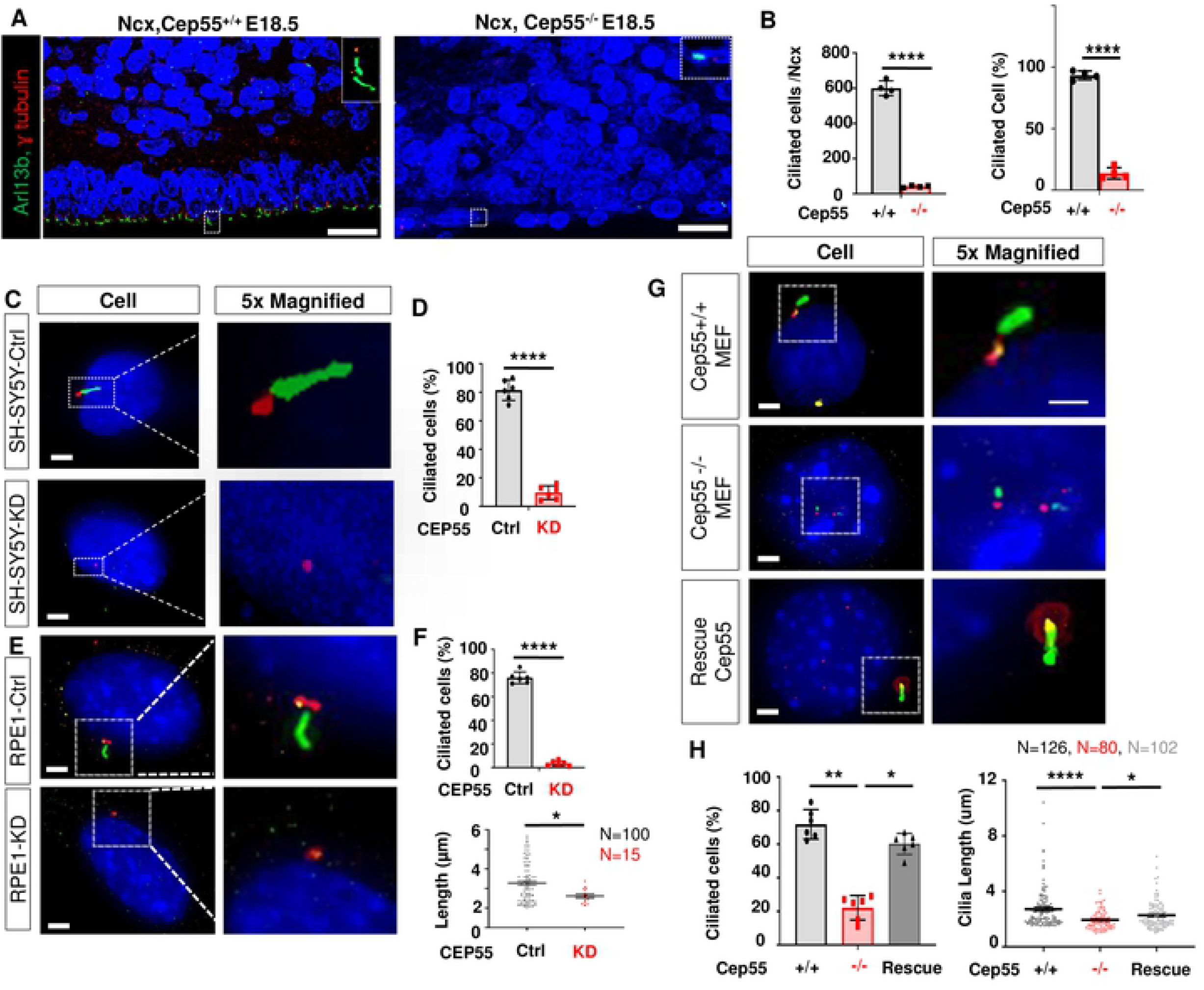
*Cep55* is localized to basal body of cilia and directly regulates its growth. **(A)** Super-resolution microscopy of *Cep55^+/+^* and *Cep55^-/-^* mouse neocortex at E18.5 immunostained for cilia (Arl13b), basal body (γ-tubulin), and DAPI, scale = 10 μm. **(B)** Percent of cilia-positive cells in Ncx at E18.5. **(C-H)** Representative images showing cilia (Arl13b), basal body (γ-tubulin), *Cep55* (yellow), and DAPI (blue), Scale=5 μm. **(C)** Representative image of cilia and basal body in control (upper) and CEP55 knockdown (lower) in SH-SY5Y cells. **(D)** Percentage of ciliated cells in Ctrl and KD SH-SY5Y. **(E)** Cilia, basal body, and Cep55 staining in RPE-1 cells transfected with a control lentiviral vector (Ctrl) (upper), RPE-1 cells with lentiviral knockdown of Cep55 (lower). **(F)** Percentage of ciliated cells in RPE-1 cells or *CEP55* knockdown (upper), Scatter plot showing cilia length in RPE-1 cells or CEP55 knockdown (lower). **(G)** Representative image of cilia, basal body and Cep55 in Cep55^+/+^ (Wt) MEFs (upper), Cep55^-/-^ (KO) MEFs (middle), Cep55^-/-^ MEFs with ectopic expression of Flag-Cep55 (Rescue) (lower). **(H)** Percentage of ciliated cells in Cep55^+/+^, Cep55^-/-^ and Cep55^-/-^ cells reconstituted with a Cep55 construct (rescue). Data represent mean ± SD of 300 cells per genotype (left); scatter plot showing cilia length (μm) in Cep55^+/+^, Cep55^-/-^ and Cep55-reconstituted MEFs. Data were measured in duplicate across two independent experiments (right). For all calculations, Student’s t-test, *P < 0.05, **P < 0.01, ***P < 0.001, ****P < 0.0001).

### CEP55 is localized to the ciliary basal body and regulates cilia growth

Finding the cilia defect in *Cep55^-/-^* brains, we aimed to investigate how Cep55 regulates ciliogenesis and whether ciliary defects could be recapitulated in an *in vitro* system to facilitate mechanistic studies. We turned our attention to a cell line of neural origin, SH-SY5Y[19], and to the hTERT RPE-1 line, which has been routinely used to examine ciliogenesis[20]. Ciliogenesis usually occurs in G_0_ and G_1_ and serum starvation is widely used to arrest cells in G1 to stimulate cilia formation. We examined the impact of C*ep*55 knockdown on cilia formation in SH-SY5Y and RPE-1 cells and found significantly decreased numbers of ciliated cells compared to the respective control; the latter also revealed a significant reduction in cilia length after *Cep55* knockdown (Fig 4C-F; fig S3C,D). Finally, to facilitate rescue studies, we isolated mouse embryonic fibroblasts (MEFs) from *Cep55^+/+^* and *Cep55^-/-^* mice to investigate possible cilia defects and perform rescue experiments with ectopic expression of CEP55 in *Cep55^-/-^* MEFs. Consistently, we found that *Cep55^-/-^* cells had a significantly reduced number of ciliated cells and shorter cilia when compared to *Cep55^+/+^* MEFs (Fig. 4G, H). In addition, a slight increase was seen in the number of *Cep55^-/-^* MEFs displaying multiple small cilia extending from the basal body (double cilia), alongside a significant proportion of cilia from KO MEFs exhibiting dissociation from the basal body (remnant cilia) (fig S3E). In order to evaluate whether defective ciliogenesis is a direct consequence of *Cep55* loss, we rescued cilia formation by ectopic overexpression of *Cep55* in *Cep55^-/-^* MEFs. We showed that ectopic *Cep55* expression (fig S3F) was able to restore cilia formation and length to levels comparable to *Cep55^+/+^* MEFs (Fig. 4G lower panel, Fig 4H). Given an apparent role for *Cep55* in ciliogenesis regulation, we also examined whether *Cep55* co-localizes with the cilia axoneme or at the base of cilia. Co-staining of Cep55 (yellow) with Arl13b (green) and gamma-tubulin (red) revealed an apparent co-localization of Cep55 with gamma-tubulin (a component of the basal body protein complex) in *Cep55^+/+^* or rescue MEFs. To investigate this, we performed super-resolution microscopy to image Cep55 (yellow) and γ-tubulin (red) across a population of Cep55-rescued MEFs (ectopic *Cep55* expression) and observed staining of both proteins at the base of cilia (fig. S3G). Together, these findings illustrate that Cep55 localizes at the base of cilia, possibly as a component of the basal body protein complex and is required for normal cilia formation.

### *Cep55^-/-^* MEFs exhibit multinucleation and cell cycle defects

Given that *Cep55^-/-^* embryos are growth restricted *in vivo*, we next sought to recapitulate this phenomenon *in vitro* to determine if *Cep55* loss causes proliferation defects in our mouse embryonic fibroblast (MEF) cell lines. We calculated cell doubling time and found significant growth defects in *Cep55^-/-^* lines (fig. S4A) compared to *Cep55^+/+^*, which was further revealed to be dose-dependent (fig. S4B), thus supporting *in vivo* growth restriction. Moreover, we were able to rescue this proliferation defect by ectopically expressed *Cep55* in *Cep55^-/-^* MEFs (fig. S4C). Overall, the *in vitro* proliferation deficiency in *Cep55^-/-^* MEFs was consistent with the observed phenotype in neuronal progenitors, where decreased proliferation was detected with Ki67 (E14.5) by immunofluorescence. Furthermore, the multinucleation seen in MEFs (fig. S4D,E) was reminiscent of the neuronal phenotype. We next performed cell cycle analysis using propidium iodide (PI)-stained cells sorted by flow cytometry. FACS analysis revealed significant differences in the cell cycle profile of *Cep55^+/+^* and *Cep55^-/-^* primary MEFs, where *Cep55^-/-^* cells showed enrichment of cells in G2 and a reduction in G1 population (fig. S4F-H). To further examine cellular division, we performed live-cell imaging of *Cep55^+/+^* and *Cep55^-/-^* cells transduced with mCherry-histone H2B by EVOS-FL time-lapse microscopy (fig. S4I). We found extended cell division (mitotic length) in *Cep55^-/-^* compared to *Cep55^+/+^* lines (fig. S4J). As expected, the *Cep55^-/-^* cells showed defective cytokinesis, taking longer to divide effectively with 17% remaining multinucleated (fig. S4K). For further characterization of additional mitotic defects, we performed high-resolution time-lapse microscopy of mCherry-histone H2B cells using Spinning Disk Confocal microscopy to quantitate mitotic defects including anaphase bridge formation, lagging chromosomes, and mitotic slippage. Although there was a trend towards an increased proportion of anaphase bridges during mitosis in *Cep55^-/-^* MEFs compared to control, this was not statistically significant (fig S4L). Moreover, we did not observe any changes in lagging chromosomes or slippage (fig S4L). Together, these results illustrate that Cep55 is important to support normal cell growth and division in particular cytokinesis in MEFs.

### Cep55 regulates Gsk3β, downstream of the Akt pathway

Cep55 has previously been shown to regulate PI3K/AKT signaling pathway in cancer cells[2,3], we initially performed signaling analysis of AKT and its downstream targets in E14.5 mouse brains as well as in MEFs by immunoblotting. The loss of Cep55 downregulated pAKT in both *Cep55^-/-^* brain tissue and MEFs, consistently (Fig. 5A, B). AKT controls steady-state levels of GSK3β through phosphorylation of residue Serine 9 (pS9-GSK3β). Inactive AKT is known to result in decreased pS9-GSK3β levels, which leads to GSK3β activation with pro-apoptotic functions[21]. In line with this, we observed that *Cep55* loss led to the inactivation of Akt (decreased pS473) and activation of GSK3β (decreased levels of pS9-GSK3β) (Fig. 5A, B). Importantly, we were able to rescue the phosphorylation of Akt and Gsk3β by ectopic expression of *Cep55* in *Cep55^-/-^* MEFs, confirming the specificity of the observed signal transduction effects (Fig 5C). Activated Gsk3β has been shown to inhibit downstream targets involved in proliferation such as β-catenin and Myc. We observed a decrease in β-catenin levels in E14.5 *Cep55^-/-^* brains (Fig 5A). Similarly, we observed reduced β-catenin and non-phospho β-catenin levels in *Cep55^-/-^* MEFs (Fig. 5B). GSK3β can also destabilize Myc by phosphorylation on Threonine 58[22]. Accordingly, we observed an increase in pT58-Myc levels and a concomitant decrease in total Myc levels in *Cep55^-/-^* MEFs as well as a trend of decreased total Myc levels in E14.5 brains when compared to *Cep55^+/+^* controls (Fig. 5A, B). Additionally, we performed IHC staining of total β-catenin and N-Myc on E14.5 brain sections from *Cep55^-/-^* and *Cep55^+/+^* embryos. These results revealed a significant decline of membranous and cytoplasmic β-catenin in *Cep55^-/-^* in the VZ compared to *Cep55^+/+^* in E14.5 brain, consistent with immunoblot analysis in the embryonic brain at this time-point (Fig 5D). Similarly, IHC on E14.5 brain sections revealed a significant reduction in N-MYC expression in *Cep55^-/-^* NPs compared to *Cep55^+/+^* controls (Fig 5E). Notably, *Cep55* loss resulted in a reduction in transcript levels of Myc in both brain samples and MEFs, consistent with reported transcriptional regulation of Myc by the Wnt/β-catenin pathway (fig S6A). Additionally, N-Myc (a member of the Myc family regulating neural cells) was shown to be reduced in E14.5 *Cep55^-/-^* brain tissue (fig S5B). Together, we conclude that *Cep55* loss potentially inhibits proliferation and survival in an AKT-dependent manner.

**Fig 5.**
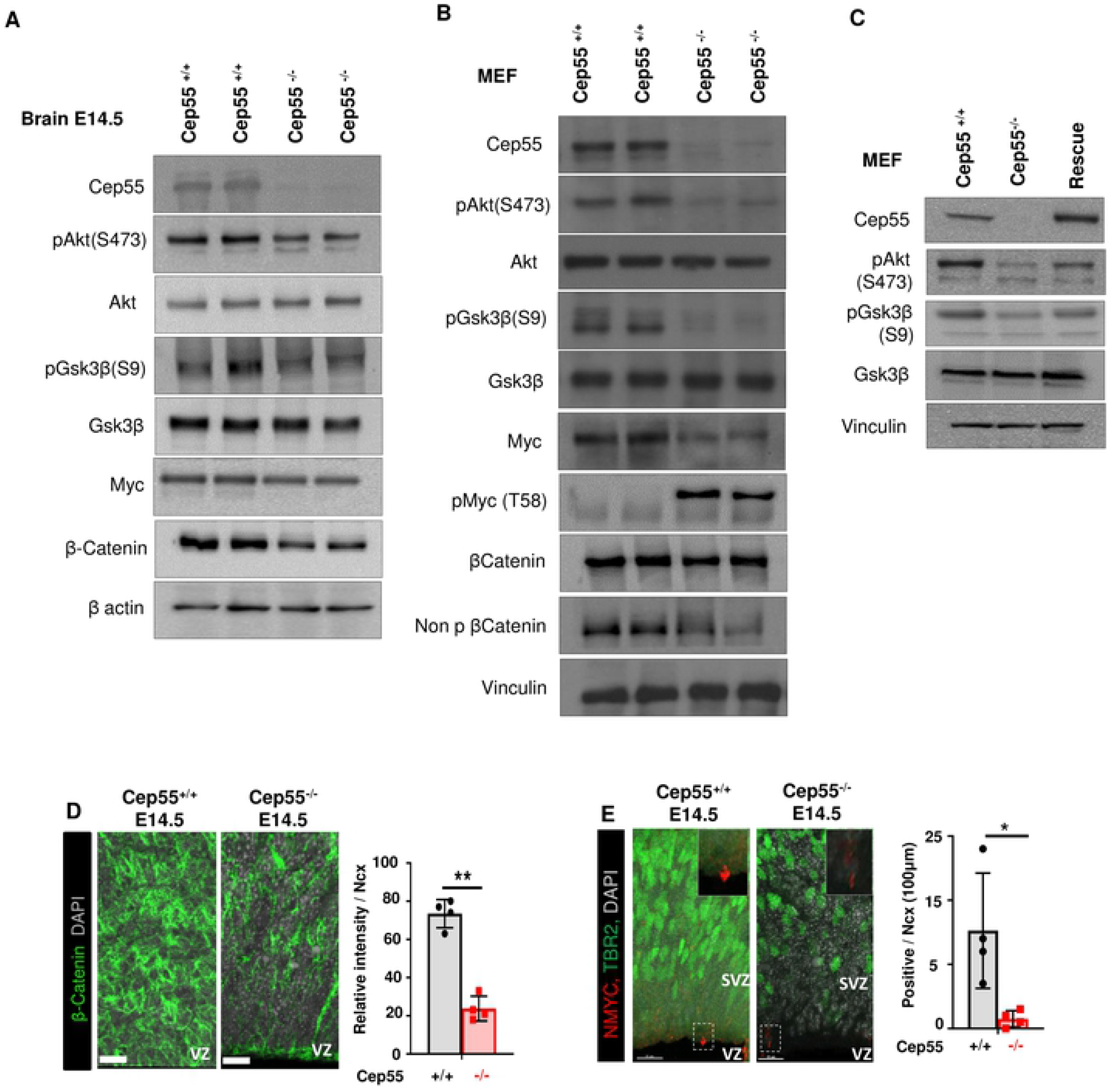
Cep55 regulates GSK3β, β-Catenin and Myc downstream of the Akt pathway. **(A-C)** Representative image of immunoblot (WB) analysis with indicated antibodies. β actin or vinculin served as a loading control. WB performed to compare **(A)** *Cep55^+/+^* and *Cep55^-/-^* mouse embryonic brain at E14.5. **(B)** *Cep55^+/+^* and *Cep55^-/-^* MEFs. **(C)** *Cep55^+/+^* (Wt), *Cep55^-/-^* (KO) and *Cep55^-/-^* MEFs with ectopic expression of Cep55 (Rescue) with indicated antibodies. **(D)** Representative images of Cep55+/+ (left) and Cep55-/- (right) neocortices stained for β-catenin and nuclei (DAPI) in a 100 μm-width box. Bar chart shows the relative intensity of β-catenin signals for Cep55+/+ and Cep55-/- neocortices; Scale= 50 μm. **(E)** Representative images of Cep55+/+ (left) and Cep55-/- (right) neocortices stained for N-Myc (red), TBR2 positive cells and nuclei (DAPI) in a 100 μm-width box. Bar chart shows the percent of N-Myc positive cells, scale= 15μm (Mean ± SD of four embryos duplicate technical repeats, Student’s t-test, *P < 0.05, **P < 0.01, ***P < 0.001, ****P < 0.0001).

### Modulating downstream effectors of the PI3K/AKT signaling pathway rescues the proliferation and cilia defects induced by *Cep55*-loss

Next, we sought to evaluate whether the reconstitution of Akt signaling or its downstream regulators would be sufficient to rescue the proliferation and ciliogenesis defects in *Cep55^-/-^* MEFs. To investigate this, we first, utilized a myristoylated form of AKT1 (myrAKT) previously described to be constitutively-active[23]. *Cep55*^-/-^ MEFs were transduced with retrovirus to express myr-AKT or empty vector (EV) control (fig S5C) and assessed for proliferation. Incucyte™ analysis revealed that myr-Akt was able to markedly increase the proliferative rate of *Cep55^-/-^* MEFs when compared to EV-transduced cells (Fig. 6A). We also sought to determine if we could rescue the proliferation and ciliogenesis defects using an inhibitor of activated GSK3β. The universal GSK3β inhibitor, CHIR99021, at low dosages (0.1μM and 1 μM) was able to partially increase proliferation in *Cep55^-/-^* MEFs (Fig. 6B), possibly through the inhibition of active GSK3β as per previous reports[24]. In contrast, in *Cep55*^+/+^ lines (similar to *Cep55* rescued lines where GSK3β is inactivated by Akt activity); GSK3β inhibition can hinder proliferation in a dose-dependent manner (fig S5D).Our findings demonstrate that Cep55, through activation of AKT and inhibition of GSK3β, can regulate proliferation.

**Fig 6.**
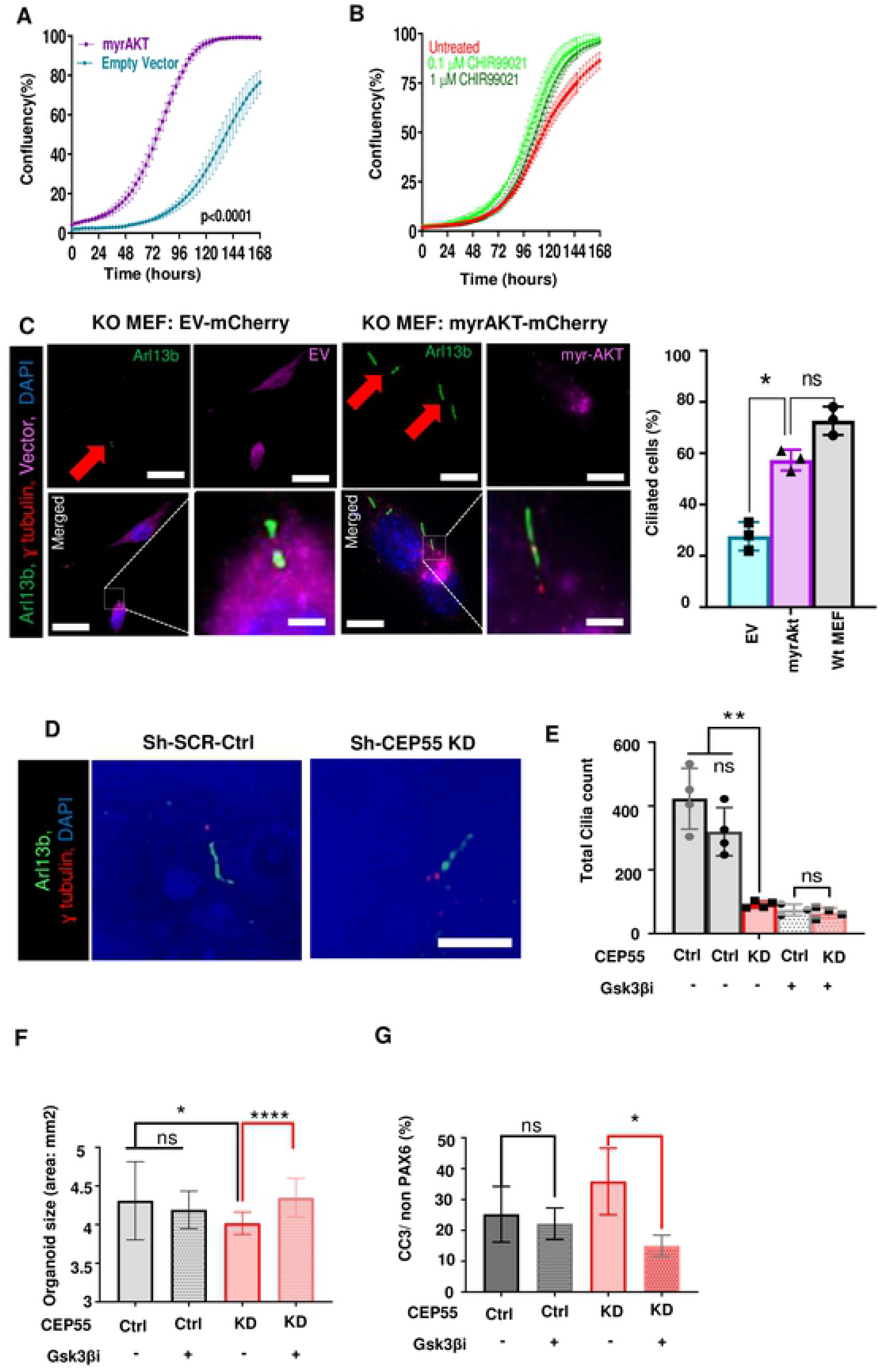
Myr-Akt and downstream effectors can rescue *Cep55* loss. Proliferation assay showing the growth of *Cep55^-/-^* (KO) MEF **(A)** Transiently transfected with EV (turquoise blue) or myr-Akt (purple) and **(B)** treated with indicated doses of GSK3β inhibitor, CHIR99021 (untreated: red, 0.1 μM inhibitor: light green, 1 μM inhibitor: dark green). Mean ± SD, average of 2 biological repeats and 3 independent experiments Student’s t-test, ****P < 0.0001. **(C)** Representative images of *Cep55^-/-^* (KO) MEFs reconstituted with EV-mCherry (left) or myrAKT-mCherry (right) immunostained for cilia (Arl13b), basal body (γ-tubulin) and nuclei (DAPI). The lower panel shows merged images with magnification of the boxed area. Scale=12 μm. Left: Percentage of ciliated cells in *Cep55*^+/+^ (Wt) and *Cep55*^-/-^ (KO) MEFs transfected with EV or myr-Akt. Mean ± SD, n=100 cells from 3 independent experiments Student’s t-test, *P < 0.05). **(D)** Representative image U6 (shSCR Ctrl) and CEP55 KD human brain organoids immunostained for cilia (Arl13b), basal body (γ-tubulin) and nuclei (DAPI), Scale=10 μm. **(E-G)** Comparison of U6 (shSCR Ctrl) and CEP55 KD untreated (DMSO) and treated with 3μM GSK3β inhibitor, CHIR99021 for **(E)** Ciliated cell counts. **(F)** The size of organoids (area). **(G)** The percent of cleaved caspase 3 in PAX6 negative cells. Data were measured across two independent experiments. n=6 organoids. Mean ± SD, Student’s t-test, **P < 0.01).

Next, we examined whether myrAKT expression was sufficient to rescue the defects in cilia formation. Strikingly, we observed that myrAKT expression in *Cep55^-/-^* MEFs but not EV expression restored the percent of ciliated cells to levels more comparable to *Cep55^+/+^* MEFs (Fig. 6C). Regarding ciliogenesis, inhibition of GSK3β in *Cep55^-/-^* MEFs at tested concentrations did not affect cilia formation significantly (fig S5E). Our analysis of cilia in CEP55 KD organoids revealed that CEP55 loss perturbed ciliogenesis (Fig. 6D). However, in accordance with MEFs data, we were unable to rescue this phenotype with GSK3β inhibitor (Fig. 6D, E). This is consistent with a previous study that showed that GSK3β inhibition alone does not modulate ciliogenesis but combined inactivation of Von Hippel-Lindau (VHL) and GSK3β leads to loss of cilia formation and maintenance[25] suggesting that GSK3β acts redundantly with VHL to regulate ciliogenesis. Strikingly, GSK3β inhibition can rescue the size of human organoids. CEP55 loss in organoids led to size (area) decrease compared to control, consistent with microcephaly seen in Cep55-null human patients and mouse model (Fig. 6F). While the CHIR99021 (Gsk3β inhibitor) treatment had no significant effect on the size of control organoids, it led to a significant increase in the size of CEP55 KD organoids (Fig. 6F). The mechanism of this rescue is through the reduction of apoptotic cells as marked by cleaved caspase 3 (Fig. 6G). Taken together, our findings demonstrate that CEP55 regulates cell survival in an AKT-dependent manner (Fig 7). Nevertheless, Cep55-dependent regulation of ciliogenesis might occur through an AKT downstream effector(s) independent from GSK3β, Overall, the phenotype in brain organoids and related apoptosis can be rescued through GSK3β inhibition.

**Fig 7.**
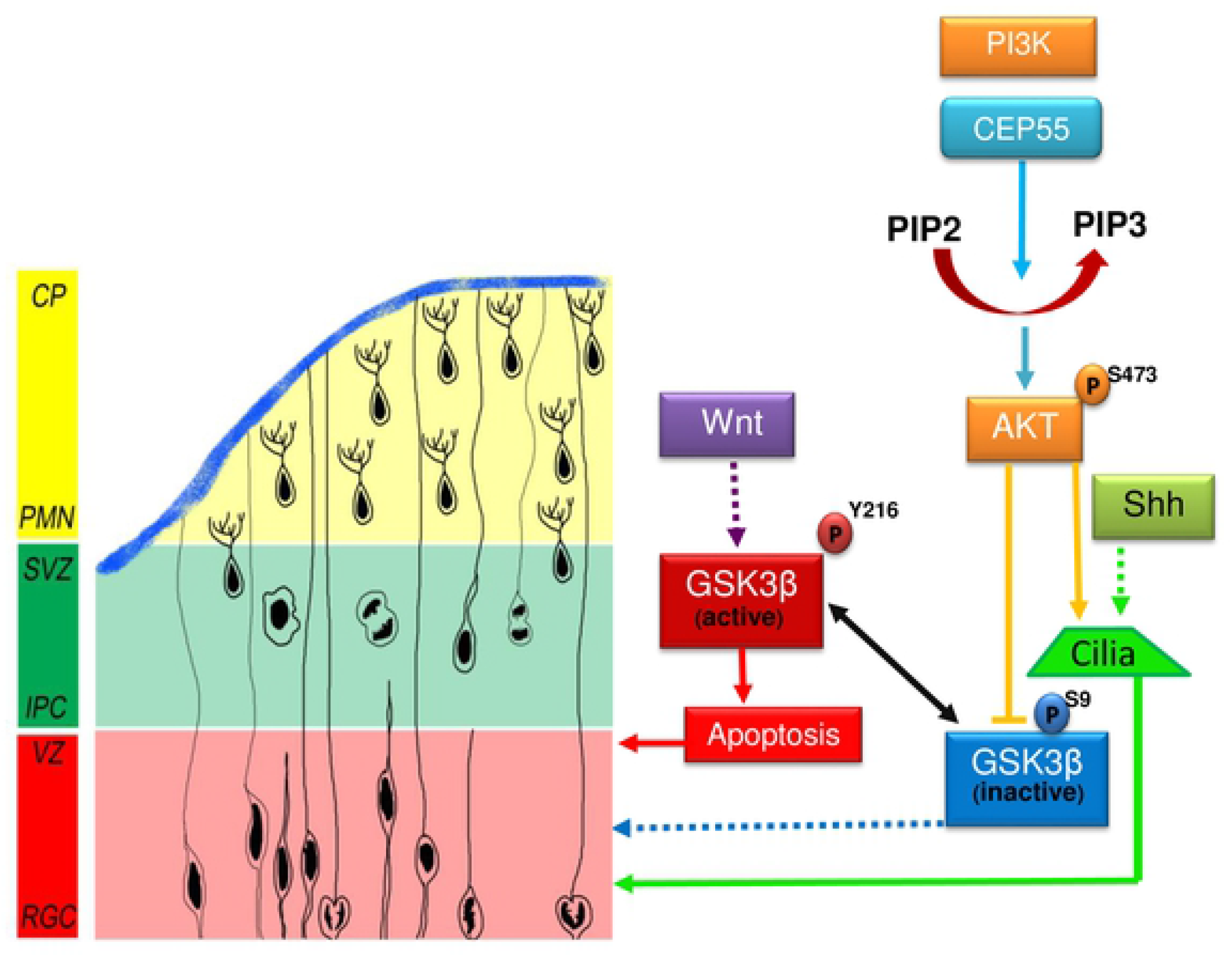
Graphical abstract. Proposed model of CEP55 regulation of RGC proliferation or apoptosis through PI3K.AKT and the downstream targets GSK3β, β-Catenin and MycCEP55 binds to the catalytic subunit of PI3K (p110) and promotes effective conversion of phosphoinositol (4,5) bisphosphate (PIP2) to PIP3 and downstream phosphorylation of AKT (S473). The active AKT inactivates *GSK3*β by phosphorylating it on S9. However, in CEP55 KO, downregulation of AKT phosphorylation leads to upregulation of the active *GSK3*β (Y216) under the regulation of Wnt signaling and can activate apoptosis. Ciliogenesis is regulated in an AKT-dependent manner in a possible crosstalk with Shh and independent of *GSK3*β.

## Discussion

CEP55 was initially described as an abscission component serving to regulate cellular segregation during cytokinesis. Later, the finding of CEP55 regulatory roles in PI3K/AKT survival signaling illustrated the importance of this protein, especially in cancer where transcriptional upregulation of *CEP55* widely contributes to cancer progression[2]. Interestingly, activating mutations in genes of PI3K pathway has been shown to cause a wide range of brain and body overgrowth disorders[26] with phenotypic severity highly dependent on the extent of activation of the pathway[27]. In contrast, the reduction in the activity of the PI3K pathway in specific organs can lead to decreased organ size[28]. Recently, four reports linked homozygous nonsense *CEP55* mutations that truncate the protein to the lethal fetal syndromes, demonstrating the importance of *CEP55* in embryogenesis and especially in neuronal development[4–6]. Surprisingly, patients compound heterozygotes for nonsense and missense or splicing mutation in Cep55 survive[29]. However, to date, the exact molecular mechanism underlying these disorders remained elusive. By simultaneously studying *Cep55^-/-^* mouse model and human cerebral organoid, our studies provided novel insights into the pathophysiological role of *Cep55* to understand the disease linked to dysregulation of this gene. The lethal phenotypes in this model, including the forebrain and hindbrain abnormalities and overall lack of proper cerebral development overlap with the human disorders. Notably, we also observed a higher proportion of multinucleated neurons in E14.5 *Cep55^-/-^* brains when compared to controls, mimicking neuron phenotype of CEP55-associated MARCH syndrome[6]. Consistently, a significant proportion of MEFs exhibited multinucleation upon both constitutive and conditional loss of *Cep55*.

The brain size of *Cep55-*deficient embryos is significantly reduced compared to controls due to hypocellularity. It is conceivable that the apoptosis seen in brain sections could progress to a major loss of cerebral hemisphere parenchyma, resulting in marked cavitation, leaving only a small amount of residual cortical tissue, and compensatory expansion of the lateral ventricles (termed hydranencephaly: seen in human with CEP55 mutation), or porencephaly, if the cystic change and parenchymal loss was less severe. For the detailed characterization of neurodevelopmental defects upon loss of *Cep55*, in this study, we have concentrated on the development of the neocortex. Analysis of the overall distribution of neurons across the neocortex revealed a decreased population of all NPs and neurons in *Cep55^-/-^* brains including RGCs, IPCs, and PMNs. Deficiency in the proliferation of *Cep55^-/-^* brain cells in the early neurodevelopment stage, as well as the significant increase of apoptosis in both NPs and PMNs, are likely to be due to the pro-survival role of Akt where its activation is significantly compromised in *Cep55^-/-^* brain cells. This finding is in line with the time-lapse data showing that cell death is not due to the mitotic catastrophe caused by aberrant cytokinesis but cells are mainly dying during interphase. Our data suggests proliferation defects associated with loss of Cep55 in both E14.5 embryo brains as well as *in vitro* in MEF models with both constitutive and conditional Cep55 loss. This proliferation defect could be rescued in *Cep55*^-/-^ MEFs by ectopic expression of *Cep55*. Overall, the *in vitro* proliferation deficiency in *Cep55*^-/-^ MEFs was consistent with the observed phenotype in neuronal progenitors, with decreased proliferation observed by Ki67 immunohistochemistry at E14.5. We also observed increased levels of cleaved caspase-3, a marker of apoptotic cell death by WB in *Cep55^-/-^* embryonic brain tissue consistent with increased apoptosis as assessed by TUNEL staining of brain sections by IHC.

In addition to the pro-survival role of Akt in regulating proliferation and apoptosis, the dysregulation of Akt[30] as well as its downstream effectors such as GSK3β [31], Myc, and β-Catenin[32] have been reported to have adverse effects on neurodevelopment, predominantly affecting proliferation. GSK3β, implicated as a master regulator of NPs, is a central mediator of a wide range of processes in neurodevelopment[33]. Our data showed reduced Akt phosphorylation and a consequent reduction in inhibitory phosphorylation on Gsk3β in the absence of Cep55 leads to Gsk3β activation and consequent proteasomal mediated degradation of its substrate, β-catenin. In the *Cep55*^-/-^ brain and MEFs we observed reduced expression of β-catenin by WB and IHC in the absence of any changes at the level of transcription. We identified Myc destabilization in MEFs and embryonic brain in protein level by WB and decrease in Myc and N-Myc transcript levels by RT-qPCR. We also validated these results by IHC analysis which revealed a decline in N-Myc protein expressed mostly in the VZ of mouse brains. Overall, reduced proliferation and increased apoptosis in NPCs upon Cep55 deletion could explain smaller brain size.

The function and regulation of Cep55 in the primary cilium have never been studied despite the fact that many centrosome localized proteins provide a template for ciliogenesis. The ciliogenesis defect observed in Cep55-depleted cells in this study represents the first example in which multiple pieces of evidence support this notion. First, *Cep55^-/-^* embryonic brain sections (E14.5) and Cep55-depleted human cerebral organoids consistently showed ciliogenesis defects in NPCs. Moreover, several other cellular models used in the study including *Cep55*^-/-^ MEFs and *Cep55*-depleted RPE-1 and SH-SY5Y cells also exhibited a primary cilium defect. This suggests that Cep55 regulates cilia across different species. This is consistent with the described association of *Cep55* with human MKS like ciliopathy syndrome[4,5]. Second, we found that *Cep55* is predominantly localized at the base of the primary cilium, therefore, ensuring appropriate cilium assembly. Third, myr-AKT overexpression is sufficient to restore the deficit in cilium length and proliferation defect in Cep55-deleted MEFs. There is emerging evidence that the cilia dysfunctions contribute to many neurogenetic disorders such as Meckel-Gruber syndrome[34]. During neurodevelopment, PI3K/Akt activation is known to mediate the downstream effects of Shh, a regulator of corticogenesis, and the main signaling regulator of cilia[35]. It has been reported that pAkt is localized to the primary cilia basal body or to a centrosome-like structure in dividing cells; consequently, Akt knockdown can suppress cilia formation[36]. We propose that defective activation of PI3K/AKT pathway in absence of *Cep55* leads to defective proliferation and survival of neurons as well as defective cilia formation. However, inhibition of activated Gsk3β observed in *Cep55*^-/-^ MEFs and human organoids as a consequence of reduced Akt activation could only rescue the phenotype through decreasing apoptosis without any impact on ciliogenesis, suggesting that other downstream effectors of AKT are involved in the regulation of ciliogenesis.

In summary, our study has used a mouse model as well as human brain organoids to identify the critical role of *Cep55* during brain development and suggests that defective PI3K/Akt pathway activation and consequently, increased apoptosis during embryogenesis could be the predominant cause of microcephaly seen in *Cep55* loss-associated genetic syndromes. In addition, we revealed an important role of Cep55 in regulating ciliogenesis in an Akt dependent manner; further studies should determine to what extent disruption of ciliogenesis contributes to complex Cep55-associated clinical phenotypes [4–6].

## Materials and Methods

### Animal husbandry and ethics statement

All experimental animals were maintained on a C57BL/6J strain. For the generation of transgenic mouse see the supplementary methods. This research was carried out in strict accordance with the Australian Code for the care and use of animals for scientific purposes. All protocols were approved by the QIMR Berghofer Medical Research Institute Animal Ethics Committee (ethics number A0707-606M).

### Immunohistochemistry (IHC) and Immunofluorescence (IF)

For IHC, tissues including embryonic brains were collected and fixed in 10% buffered formalin fixative, 4% Paraformaldehyde or Bouin’s solution (Sigma-Aldrich), embedded in paraffin blocks, and 5-10 μm-thick sections were stained with Haematoxylin and eosin or with indicated antibodies. Slides were examined by an independent veterinary pathologist. Immunohistochemistry staining was performed following standard procedures. Stained slides were scanned on an Aperio ScanScope Fl Slide Scanner (Leica) or imaged on a Zeiss 780-NLO - Confocal microscope (Zeiss, Jena Germany) and images analyzed using Imaris or ImageJ software. (See the supplementary methods). Immunofluorescence (IF) assays were performed as previously described^1^ (See the extended methods).

### Cerebral brain organoid differentiation

Cerebral brain organoids were generated from HES3 pluripotent cultures were single-cell dissociated using Accutase as per manufacturer’s instructions (Life Technologies). (For detailed procedure see the supplementary methods).

### Cell proliferation assay

Cells were seeded at a density of 5×10^3^ or 10^4^ cells per well in duplicate, and growth assessed using an IncuCyte^®^ S3 Live-Cell Analysis system (Essen BioSciences Inc, USA) Where treatments were performed, drugs were added the day following cell seeding.

### Immunoblotting

Western blotting was performed as previously described[3]. Protein detection was performed using Super Signal chemiluminescent ECL-plus (PerkinElmer, Waltham, MA, USA) on a BioRad Gel doc (Bio Rad ChemiDoc Touch, USA). More details are available in supplementary methods (See the extended methods).

### Quantitative real-time PCR

Reverse Transcription was performed using the SuperScript First-Strand Synthesis System for RT-PCR. This cDNA was then used as the template for real-time PCR with gene-specific primers as outlined in supplementary methods.

### Statistical analysis

Two-tailed unpaired or paired Student’s t-test, one-way or two-way ANOVA with post hoc Bonferroni, log-rank testing were performed as indicated using Prism v8.0 (Graph Pad Software, La Jolla, CA, USA) and the P-values were calculated as indicated in the figure legends. Mean and standard error of mean (Mean ± SD) are used to describe the variability within the sample in our analysis. ns; P > 0.05; *P ≤ 0.05; **P ≤ 0.01, ***P ≤ 0.001, and ****P ≤ 0.0001.

## Acknowledgments

We acknowledge the International Knockout Mouse Consortium for providing Cep55 targeted embryonic Stem Cell Clones (Project ID: 93490), Australian Phenomics Facility (Melbourne) for the generation of knockout mice and histopathology analysis of embryos, and QIMRB for Animal Facilities, Histology, Flow Cytometry and Microscopy Facilities, as well as Nigel Waterhouse and Tam Hong Nguyen for their help. We thank Mathew Jones (Diamantina Institute, UQ) for providing RPE1 cells, Rick Pearson (Peter Mac Callum Cancer Institute, Melbourne) for myrAKT, and Jill Larson (QIMR Berghofer) for Lenti-Cep55 expression construct. We would like to appreciate Khanna laboratory members for critical discussions and technical assistance.

## Author Contributions

Conceptualization: BR, ALB, DS, MK, BS, MP and KK; Investigation: BR, BS (Histology), and PF (Organoid); bioinformatics: BR; Supervisory team: ALB, DN, JH, AL and KK; Pathology: JF, PB, MP, APN; data analysis: BR, BS, DS; AB (Mitosis); Writing the original draft, BR, ALB and KK; review and editing: BR, ALB, RA, PF, MP, JF, JH, DN and KK; All authors read and approved the final manuscript.

## Supplementary Information

### Supplementary Tables

**Table S1.** Proportion of observed and expected offspring from *Cep55^+/-^* x *Cep55^+/-^* intercrosses at time of birth (P=0).

| P0 | WT | HET | KO | Total |
| --- | --- | --- | --- | --- |
| **Observed** | 34 | 43 | 0 | 77 |
| **%** | 44.15% | 55.85% | 0% |  |
| **Expected** | 19.25 | 38.5 | 19.25 | 77 |
| **%** | 25% | 50% | 25% | 1 |

**Table S2.** Number and percentage of offspring at indicated stages of gestation from *Cep55^+/-^* X *Cep55^+/-^* intercrosses.

| Embryonic Day | Attempts | WT | HET | KO | Total |
| --- | --- | --- | --- | --- | --- |
| **E11.5** | 1 | 6 | 1 | 3 | 10 |
| % |  | 60% | 10% | 30% | 100% |
| **E13.5** | 7 | 18 | 20 | 16 | 54 |
| % |  | 33.3% | 37% | 29.6% | 100% |
| **E14.5** | 4 | 5 | 17 | 6 | 28 |
| % |  | 17.9% | 60.7% | 21.4% | 100% |
| **E15.5** | 1 | 3 | 5 | 3 | 11 |
| % |  | 27.3% | 45.5% | 27.3% | 100% |
| **E16.5** | 3 | 7 | 14 | 4 | 25 |
| % |  | 28% | 56% | 16% | 100% |
| **E18.5** | 4 | 11 | 14 | 12 | 37 |
| % |  | 29.7% | 37.8% | 32.4% | 100% |
| **P0** | 2 | 3 | 5 | 0 | 8 |
| % |  | 37.5% | 62.5% | 0% | 100% |
| **Total** | 22 | 53 | 76 | 44 | 173 |
| % |  | 30.6% | 43.9% | 25.5% | 100% |

### Supplementary figure legends

**Fig S1.**

**(A)** Schematic representation showing Wt (wild type), transgenic (gene trapped), floxed and knockout alleles of murine *Cep55*, blue arrows indicate genotyping primers. **(B)** PCR genotyping showing *Cep55^+/+^, Cep55^+/-^* and *Cep55^-/-^* genotypes. **(C)** mRNA expression of *Cep55* in *Cep55^+/+^, Cep55^+/-^* and *Cep55^-/-^* E14.5 mouse heads. *ACTB* was used as a housekeeping gene for normalization. Data represent the mean ± SD, n = 2 mice per genotype, 3 independent experiments, Student’s t-test *P < 0.05, **P < 0.01, ***P < 0.001, ****P < 0.0001). **(D)** Immunoblot analysis of Cep55 protein expression from *Cep55^+/+^, Cep55^+/-^* and *Cep55^-/-^* E14.5 mouse heads. β-actin was used as a loading control. **(E)** Comparison of organ volumes of 8-week-old *Cep55^+/+^* and *Cep55^+/-^* mice. Brain and thymus size are slightly smaller in *Cep55*^+/-^ (Het) mice, n=2 per group. **(F)** Mean body weights of *Cep55*^+/+^ and *Cep55*^+/-^ offspring measured at the indicated time points until 20 weeks. n= 6-9 mice per group. **(G)** *Cep55* expression in the single-cell transcriptomic analysis of mouse neocortical development visualized based on the available data at Zylka lab dataset. The highest expression is seen in radial-glial cells (RG2) at embryonic day 14. **(H)** β-galactosidase staining of coronal sections of *Cep55*+/-mouse embryonic brain at the indicated time points. Dotted black box indicates the magnified area shown on right, Scale=100 μm.

**Fig S2.**

**(A)** Cerebellar hypoplasia in *Cep55^-/-^* (lower panel) compared to *Cep55^+/+^* (upper panel) brain sections. Compared to the *Cep55^+/+^*, there is a marked reduction in thickness of the external granular layer (EGL) in a *Cep55^-/-^* brain. The cerebellar cortical neuronal population in *Cep55^-/-^* is deficient and disorganized, higher power views of both cerebellar cortices are also shown (right). The olfactory bulb is also neuron-deficient and disorganized in a *Cep55^-/-^* mouse compared to a *Cep55^+/+^* mouse (left). Scale = 60μm. **(B)** Comparison of cerebral hemisphere (neocortex (NCx), germinal epithelium (GE) and lateral ventricles) from Cep55^+/+^ (upper) and Cep55^-/-^ (lower) E18.5 embryos. Red arrows indicate structural dilation, distortion and disorganization, and necrotic area with neural tissue loss, scale= 200μm. Middle: magnification of boxed area showing depletion of subependymal germinal neuroblasts in Cep55^-/-^, scale= 50μm. Right: magnification of boxed area showing neocortical neuronal depletion in cerebral hemispheres and reduction of cortical neuronal population in Cep55^-/-^. Red arrow identifies multinucleated neurons. Scale = 20μm. **(C)** Hematoxylin and eosin staining of E18.5 Cep55^-/-^ cerebral cortex. Upper: neocortical hypoplasia/dysplasia. Diminished and disorganized neurons with an area of parenchymal necrosis (N, black arrow) and neural tissue loss. Phagocytosed neuronal cellular debris is arrowed and magnified. Scale=120μm. Lower: numerous bi-nucleated neurons (red arrows), scale=180μm. **(D)** Representative image of NeuN (brown) and Eosin (pink) immunohistochemical staining of E18.5 sections from Cep55^+/+^ (left) and Cep55^-/-^ (right) E18.5 embryonic brain sections showing multinucleation, scale = 50μm. **(E)** Graphical representation of percentage of total cells showing multinucleation. **(F)** Immunoblotting showing cleaved caspase 3 expression in *Cep55^+/+^* and *Cep55^-/-^* MEFs. β actin was used as a loading control.

**Fig S3.**

**(A)** Representative image of E14.5 mouse neocortex immunostained for cilia (Arl13b), basal body (γ-tubulin), and DAPI. *Cep55^+/+^* (upper) and *Cep55^-/-^* (lower), Arl13b channel and merged image are shown and Cep55 signals could not be detected in Cep55^-/-^ (left), Bar chart shows cilia-positive cells in Ncx in the 100 μm-width box at E14.5, cilia counts normalized to total cell (DAPI) number (lower) at E14.5 (right). **(B)** Cilia-positive cells in Ncx in the 100 μm-width box at E18.5, quantification of ciliated IPCs (left) and RGCs (right) in the neocortex, expressed as a ratio. Cell numbers were obtained from data shown in FigureS3A and Fig 2A,B (Mean ± SD of four embryos measured in duplicate, Student’s t-test, *P < 0.05, **P < 0.01, ***P < 0.001, ****P < 0.0001). **(C)** Representative images of RPE-1 cells transiently transfected with si-Scramble (left panel) or siRNA against *CEP55* for 48 h (right panel) showing cilia (Arl13b) and nuclei (DAPI). Bar chart shows a comparison of percentage of ciliated cells. **(D)** Immunoblot of *CEP55* expression in Ctrl (Empty vector), or CEP55-depleted (shRNA CEP55) RPE-1 cells. Vinculin was used as a loading control. **(E)** Representative images of different phenotypes of cilia in *Cep55^+/+^* and *Cep55^-/-^* MEFs (shortened cilia, double cilia and remnant cilia). Bar charts show percentage ciliated cells, cilia number and percent of cells with remnant cilia or double cilia. (Mean ± SD, n=300 cilia per group of 2 independent experiments. Student’s t-test, *P < 0.05, **P < 0.01, ***P < 0.001, ****P < 0.0001), scale= 10μm. **(F)** Immunoblotting showing *Cep55* expression in *Cep55^+/+^, Cep55^-/-^* MEFs without or with reconstituted Cep55 (rescue). Vinculin was used as a loading control. **(G)** Representative images of individual channels showing cilia (Arl13b,green), basal body (γ-tubulin,red), DAPI (blue) as well as cilia (green) and *Cep55* (yellow) and DAPI (blue) showing the co-localization of *Cep55* and γ-tubulin at the base of cilia. Scale=5 μm.

**Fig S4.**

**(A)** Doubling time of *Cep55^+/+^* and *Cep55^-/-^* MEFs (Mean ± SD, n=2 biological repeats and 3 independent experiments Student’s t-test, ****P < 0.0001). **(B-C)** Proliferation of **(B)** *Cep55^+/+^, Cep55^+/-^* and *Cep55^-/-^* MEFs and **(C)** *Cep55^+/+^* (Wt), *Cep55^-/-^* (KO) and Cep55-reconstituted (Rescue) MEFs (Mean ± SEM, average of 2 biological repeats and 2 independent experiments Student’s t-test, ****P < 0.0001), measured using IncuCyte, Corresponding immunoblotting for *Cep55* expression is shown below each graph. Vinculin was used as a loading control. **(D)** Representative images of individual channels showing α-tubulin (cytoskeleton), Cep55, and nuclei (DAPI) in *Cep55^+/+^* (left) and *Cep55^-/-^* (right) MEFs. **(E)** Bar chart showing percent of multinucleated cells in constitutive MEF (*Cep55*^+/+^ (wt), *Cep55^+/-^* (Het) and *Cep55+/-*(*KO*)), (Mean ± SD, n=300 cells counted from 2 biological repeats and 3 independent experiments Student’s t-test, *P < 0.05, **P < 0.01, ***P < 0.001). **(F-G)** Modfit histogram of cell cycle analysis by FACS showing cell cycle distribution of Cep55^+/+^ **(F)** and Cep55^-/-^ **(G)** MEFs. **(H)** Graph showing percent of cells in G1, S and G2 for each genotype. Data represent mean ± SD of two lines per genotype, measured in duplicate across three independent experiments. **(I)** Representative images from time-lapse microscopy of *Cep55*^+/+^ (upper panel) and *Cep55*^-/-^ (lower panel) MEFs transfected with mCherry-histone H2B showing different phases of mitosis and cytokinesis. **(J)** Dot plot showing the time cells take to complete the mitosis (left), the stacked bar chart showing the average time to complete cell division (right) for *Cep55^+/+^* and *Cep55^-/-^* MEFs. **(K)** Column chart showing the percentage of cells with cytokinesis failure (multinucleated cells) or success (single cells) for *Cep55*^+/+^ and *Cep55*^-/-^, (Mean ± SD, n=10-25 cells counted from 3 technical repeats Student’s t-test, **P < 0.01). **(L)** The stacked bar chart represents a comparison of percentages of different mitotic phenotypes of MEFs transfected with Cherry-histone H2B for *Cep55^+/+^* (left) and *Cep55^-/-^* (right), based on images of the cell captured by time-lapse microscopy (Spinning disk confocal microscopy). (Mean ± SD, n=55-67 cells counted from 3 technical repeats Student’s t-test, **P < 0.01).

**Fig S5.**

**(A-B)** Fold change of mRNA expression of the indicated transcripts for Cep55^+/+^ and Cep55^-/-^ E14.5 brain extracts **(A)** and MEFs **(B)**. **(C)** Immunoblot showing expression of Akt in EV and myrAkt transfected Cep55^-/-^ MEF. Vinculin was used as a loading control. Proliferation assay showing growth of **(D)** *Cep55^+/+^* (Wt, left), and Flag-*Cep55* reconstituted *Cep55*^-/-^ MEFs (Rescue, right) treated with indicated doses of GSK3β inhibitor, CHIR99021 (untreated: red, 0.1 μM inhibitor: light green, 1 μM inhibitor: dark green), (Mean ± SD, average of 2 biological repeats and 2 independent experiments Student’s t-test, ****P < 0.0001). **(E)** Representative images of *Cep55*^+/+^(left) and *Cep55*^-/-^ (right) MEFs untreated (upper) and treated (lower) with 1μM of GSK3β inhibitor, CHIR99021. Bar chart shows the percentage of ciliated cells in *Cep55^+/+^* and *Cep55^-/-^* MEFs untreated and treated with 1μM of GSK3β inhibitor, CHIR99021, n=100.

## Supplementary Materials

### Generation of constitutive and conditional Cep55 knockout mice

Mice were housed at the QIMR Berghofer Medical Research Animal Facility in OptiMICE^®^ caging (Centennial, Colorado, USA) at 25°C with a 12-hour light-dark cycle. *Cep55* floxed ES cells were purchased from the International Knockout Mouse Consortium (Exon 6 of Cep55 was trapped, IKMC Project ID:93490) and heterozygous *Cep55* targeted (*Cep55*) mice were generated by the Australian Phenomics Network (APN) facility, where the targeted allele acts as a gene-trap to form a non-functional (KO) allele. The knockout-first allele used in the targeting strategy is amenable to the generation of a floxed allele via FLP recombinase breeding, allowing the generation of conditional knockout mice. To obtain *Cep55* cKO mice, *Cep55^Tg^*^/+^ heterozygous mice were crossed with Flpe mice to remove the neo cassette and backcrossed to wild-type to remove the Flpe transgene. Heterozygous *Cep55^Fl^*^/+^ mice were intercrossed to obtain Cep55^Fl/Fl^ offspring, and crossed to RosaCre^ERT2^ transgenic mice to obtain RosaCre^ERT2+^; *Cep55*^Fl/+^ mice.

### Genotype analysis

Genotyping was performed using genomic DNA extracted from mouse ear using the QuickExtract™ DNA Extraction Solution (Lucigen, USA) according to the manufacturer’s protocol. Wild-type and *Cep55* transgenic alleles were genotyped using a 3-primer PCR with a common forward primer (P1) and two different reverse primers (P2 and P3) to differentiate between different allele forms. Primer sequences were as follows: *Cep55* P1(TGGGTCTTTAACTCATGGTC), *Cep55* P2(AGGAGTGAAAAGTCCTCACA), *Cep55* P3(GTACCGCGTCGAGAAGTT), FLPe Fwd (GTGGATCGATCCTACCCCTTGCG), FLPe Rvs(GGTCCAACTGCAGCCCAAGCTTCC). Cre F (TGTGGACAGAGGAGCCATAAC), Cre R (CATCACTCGTTGCATCGACC).

### qRT PCR

The cDNA was then used as the template for real-time PCR with gene-specific primers. A control reaction was performed without reverse transcriptase to ensure no genomic DNA had contaminated the samples, as well as a no-template DNA control. The qRT-PCR was performed in 96-well plate format using a SYBR Green master-mix (Roche Applied Science, Basel, Switzerland) with a CFX96 Touch Real-Time PCR Detection System (Bio-Rad Laboratories, US). The total volume of each reaction was 8 μL including 4 μL of Sybr green, 1 μL of each primer (1 picomole of each), 1μL of cDNA (10ng) and 2 μL of sterile water. The specificity of qRT PCR amplification was examined by checking the melting curves and running each sample on a 2 % agarose gel. The results were analyzed by the ΔΔCt method. Actin was used as a housekeeping gene.

### qRT PCR primers

*Myc* (CGGACACACAACGTCTTGGAA / AGGATGTAGGCGGTGGCTTTT), *Mycn* (CCTCCGGAGAGGATACCTTG / TCTCTACGGTGACCACATCG), *Cep55* (CCTAGTAGCTCCAAGTCAGAC / ACCTTAGGTGGTCTTTGAGTC)

### Organ/embryo isolation

Mouse organs were isolated using the Nikon SMZ45 stereo dissecting microscope (Nikon Inc, Tokyo, Japan). The isolated organs were washed in ice-cold Phosphate Buffered Saline (PBS). For protein or mRNA extraction, the samples were snap-frozen on dry ice. For histology staining, tissues were fixed in either the Bouin solution (pathology investigation) or 4% PFA (immunohistochemical staining) for 24-48 hours.

### MEF establishment

MEFs were isolated from E13.5 embryos from *Cep55^+/-^* inter-crosses for the constitutive MEF and *Cep55^fl/+^* Cre^ERT2^ X *Cep55^fl/fl^* crosses for conditional MEFs. Embryos were dissected into ice-cold sterile PBS, followed by removal of the internal viscera and head for genotyping. The remaining tissue was incubated in trypsin-EDTA (Sigma Aldrich^®^, St Louis, USA) and disaggregated by mechanical shearing using a sterile scalpel blade. The dispersed tissues were further homogenized by trituration and transferred into 25cm^2^ flasks (Corning^®^) and allowed to adhere overnight. Primary MEFs were maintained in Dulbecco’s Modified Eagle’s Medium (DMEM) (Life Technologies TM, Carlsbad, CA, USA) containing 20% Fetal Bovine Serum (SAFC BiosciencesTM, Lenexa, USA) 1% penicillin-streptomycin (Life Technologies) and 1% Amphotericin B. Primary MEFs prior to passage 5 were used for experiments as indicated. Retroviral SV40 transfection was used for the immortalization of MEFs.

### Cell culture

Mouse embryonic fibroblasts (MEFs) were generated as per extended methods. Retinal Pigment Epithelium (RPE-1) cells were obtained from the Diamantina Institute, UQ. Human neuroblastoma (SH-SY5Y) cell line was cultured in a 1:1 mix of DMEM and F-12 supplemented with NEAA (1%, non-essential amino acids), FCS (10%) and pen/strep (100 U/ml). Pluripotent stem cell line HES3 (WiCell) was maintained in mTeSR1 media (StemCell Technologies) and passaged every 4 days using ReLeSRTM as per manufacturer’s instructions (StemCell Technologies) and reseeded at 12,000 cells per cm2 onto T25 cell culture flasks coated with Matrigel (Corning). All the cell lines were routinely tested for Mycoplasma infection by Scientific Services at QIMR Berghofer Medical Research Institute.

### Cerebral brain organoid differentiation

Single-cell suspensions were counted and seeded at 10^4^ cells per well of U bottom 96 well ultra-low attachment plates (Corning) in mTeSR1 media supplemented with 10μM ROCK inhibitor and centrifuged at 300g for 3 minutes to allow for initial aggregation. The following day, media in each well was replaced with knockout serum replacement (KSR) media, with ingredients from Life Technologies (USA) consisting of DMEM/F12, 20% KSR, 1x Penicillin/Streptomycin, 1x Glutamax, 1x Non-essential amino acids and 0.1mM β-mercaptoethanol. KSR media was supplemented with 2μM Dorsomorphin and A83-01 (Sigma-Aldrich) and changed daily for the first 5 days of induction. Between days 5 and 6 of induction, media was changed at a 1:1 ratio with neural induction media consisting of DMEM/F12, 1x N2, 1x Glutamax, 1x Non-essential amino acids, 1x Penicillin/Streptomycin and 10μg/mL Heparin (Stem Cell Technologies) supplemented with 1μM CHIR99021 and SB-431542 (Sigma-Aldrich). At day 7, media changes consisted only of neural induction media supplemented with 1μM CHIR99021 and SB-431542 until day 14, at which point cultures were changed to neural differentiation media consisting of 1:1 base media of DMEM/F12 and Neurobasal, 1x Glutamax, 1x Non-essential amino acids, 1x N2 and B27 (with vitamin A) supplements, 1x Penicillin/Streptomycin, 0.05mM β-mercaptoethanol and 2.5μg/ml insulin (Life Technologies).

### Adenoviral shRNA infections

Cerebral brain organoids were infected with adenoviral shRNA viruses as per manufacturer’s instructions (Vector Biolabs). Two control and CEP55 adenoviral shRNAs were used: scrambled control Ad-U6-RNAi (cat# 1640), CMV driven Ad-GFP control (cat# 1060) and human CEP55 shRNA silencing adenovirus (cat# shADV-204994). Day 16 cerebral organoids were infected with the control and CEP55 adenoviral shRNAs at a MOI of 10. Organoids were harvested 24- and 48-hours post-infection to characterize knockdown of CEP55. (For transfection of other genes and transduction see the extended methods).

### Doubling time assay

MEFs were plated in a 10 cm petri dish, at a density of 10^5^ cells per well, in triplicate for each genotype. Every second day, cells were collected, and the overall cell number assessed using a Countess^®^ automated cell counter (Life Technologies) for a total of 6 days.

### Cell cycle analysis

Cells were plated in a 6-well plate in duplicate at a density of 10^5^ cells per well and harvested in trypsin-EDTA (Sigma Aldrich^®^, St Louis, USA) at indicated time points and fixed in ice-cold Ethanol for 24h. Cells were stained in 1mg/mL of propidium iodide (Sigma Aldrich^®^) and 15mg/mL RNAse A) at 37°C in the dark. DNA content was assessed using a FACScanto II flow cytometry (BD Biosciences, Mountain View, CA). The proportion of cells in G_0_/G_1_, S phase and G2/M were quantified using ModFit LT™ 4.0 software (Verity Software House, Topsham, ME, USA).

### Live-cell imaging and microscopy

Live-cell imaging was performed on an EVOS Fl Auto (ThermoScientific) or Spinning disk confocal (Andor) microscope using MetaMorph^®^ Microscopy automation and image analysis software. Images were analyzed using analySIS LS Research, version 2.2 (Applied Precision).

### Immunofluorescence

For immunofluorescence (IF) assays, cells were counted and seeded at 5×10^4^ cells on sterile glass coverslips. For assessing cilia, MEFs were serum-starved for 48 h prior to analysis. Where indicated, drugs were added 12-24 h prior to fixation. Cells were fixed in 4% PFA (Sigma Aldrich^®^, St Louis, USA) or ice-cold Methanol (100%) for the centrosomal protein in PBS for 20 minutes at RT and permeabilized in 0.1% TritonX-100 (Sigma Aldrich^®^) for 10 minutes or 90 seconds (cilia experiments) and blocked in 3% or 1% (cilia experiments) bovine serum albumin (BSA; Sigma Aldrich^®^) in PBS for 1 hour in a humidified chamber at RT. Coverslips were washed and incubated with Alexa-fluor-conjugated secondary antibodies (Sigma Aldrich^®^) diluted in 3 or 1% BSA (1:1000) for 30 minutes at 37°C in a humidified chamber in the dark. Coverslips were mounted using Prolong^®^ gold anti-fade mounting medium (Life TechnologyTM). Imaging was performed on a DeltaVision personal DV deconvolution microscope and DeltaVision™ Ultra (super-resolution) (Applied Precision, GE Healthcare, Issaquah, WA) and analyzed using the GE DeltaVision software package. Automated counting was performed using script modules of Fiji ImageJ software (Java3D, Minnesota, USA).

### Whole-mount immunostaining

Brain organoids were stained following a previously published protocol^38^. Briefly, organoids were fixed in 1% paraformaldehyde solution overnight at 4°C. After washing, organoids were incubated for 4 hours at room temperature in a blocking buffer consisting of 5% FBS and 0.2% TritonX in PBS. Organoids were incubated with primary antibodies (extended methods) in blocking buffer overnight at 4°C, followed by washing in blocking buffer and subsequently incubated with secondary antibodies and Hoechst (1:1000) overnight at 4°C. Organoids were again washed twice in blocking buffer at 4°C and subsequently mounted onto microscope slides using Prolong glass antifade mountant (Life technologies). Live imaging was carried out using an Andor WD Revolution spinning disk microscope to assess an increase in integrated viral GFP. Immunostained samples were imaged using a Zeiss 780-NLO confocal microscope. Four random fields of view were imaged per organoid and manually quantified using Fiji software.

### β-Galactosidase staining

Detection of β-Galactosidase Activity using LacZ reporter and X-gal Staining was performed as described by (Burn, 2012). X-gal (5-Bromo-4-chloro-3-indoxyl-beta-D-galactopyranoside, GoldBio) was used to detect reporter gene expression marked by a dark blue stain. Briefly, whole embryos/organs were dissected and fixed (4% PFA for 30 minutes) following by washing (three times with wash buffer (0.02% NP-40, 0.01% deoxycholate in PBS) and chromogenic staining with staining solution (5 mM K3Fe(CN)6, 5 mM K4Fe(CN)6, 0.02% NP-40, 0.01% deoxycholate, 2 mM MgCl2, 5 mM EGTA, 1 mg/mL X-gal in PBS) in the dark at 37°C overnight.

### Neurohistopathological analysis and immunohistochemistry staining

For histopathologic investigation with hematoxylin and eosin (H&E), tissues were collected and fixed Bouin’s solution (Sigma-Aldrich, USA) for 48 h and embedded in paraffin blocks. 5 μm–thick sections were prepared for H&E staining with a Leica Autostainer XL. For periodic acid–Schiff (PAS) staining, whole testes were removed from male mice and fixed in 4% PFA for 24-48 h. Tissues were embedded in paraffin and Wax embedded tissues were sectioned at 5-10 μm and mounted onto Superfrost Plus slides (Thermo Fisher Scientific) using the Sakura Tissue-Tek^®^ TEC™ (Sakura Finetek, Tokyo, Japan). The slides were then later dewaxed and rehydrated by standard protocols.

Antigen retrieval was performed with 10 mM Sodium citrate buffer pH=6.0 using a Decloaking Chamber ™ NxGen (Biocare Medical, USA) for 15 min at 95°C. Sections were permeabilized and blocked in blocking buffer at RT for at least 1 h (20% FBS / 2% BSA / 0.2% TritonX in PBS. Primary antibodies were diluted in blocking buffer and incubated at 4°C overnight in a humidified chamber. Alexa-fluor-conjugated (Life Technology) secondary antibodies were incubated at RT for 3 hrs in a humidified chamber. Slides were mounted with Vectashield (Vector Laboratories, Burlingame, CA, USA) followed by cover-slipping using a Leica CV5030 (Leica Biosystems, Wetzlar, Germany) glass coverslipper and Shandon Consul-Mount mounting media (Life TechnologyTM). Slides were scanned with Aperio^®^ Scanscope^®^ FL/XT (Aperio^®^, Vista, USA) using 20X or 40X magnification and imaged with an LSM780 confocal microscope (Zeiss, Jena Germany) before analyzing with Image Scope software (Leica Biosystems, Buffalo Grove, IL, USA). Nuclei count v9 algorithm or Imaris (Bitplane Scientific Software, Belfast, United Kingdom) was used to score immunopositive cells.

### Immunostaining antibodies

Immunostaining was performed with the following primary antibodies: Ki67 1:500 (rabbit, NCL-ki67p; Novacastra, Wetzlar, Germany); mouse anti γ-tubulin (1:400, T5326 Sigma), rabbit anti γ-tubulin (1:400, T5192 Sigma), rabbit anti Arl13b (1:300, 17711-1-AP Proteintech), mouse anti α-tubulin (1:300, T5168 Sigma), mouse anti Arl13b (1:300, 75287, Antibodies Inc.), rabbit anti TBR1 (1:200, ab31940 Abcam), Monoclonal Anti-Acetylated Tubulin antibody produced in mouse clone 6-11B-1(Sigma Aldrich^®^), rat anti TBR2 488 (1:200, 53-4875-80 eBioscience), rabbit anti phospho-histone3 (1:300, ab47297 Abcam), mouse anti PAX6 (1:200, DSBH), Cep55 (1:500; sc-374051Santa Cruz biotechnology), Pericentrin (1:1000; Covance, PRB-432C), β-Catenin (1:1000; Cell Signaling Technology, 9582), Cleaved Caspase-3 (1:500; 9664 Cell Signaling Technology), Tuj1 (TU20) (1:200, 4466s Cell Signaling Technology), GFP (1:500, AB290 Abcam), DAPI was used for the nuclear staining (D9564; MilliporeSigma). ApopTag staining was performed with an ApopTag peroxidase in situ apoptosis detection kit (S7100; MilliporeSigma, Billerica, MA, USA).

### Gene transduction and transfection

For the generation of stable and constitutive cell lines with overexpression or knockdown of Cep55, we used Flag-Cep55 cloned into the pLenti PGK Hygro Dest vector (addgene#19066), or mouse small-hairpin RNAs (shRNAs) in the pLKO plasmid (Sigma Aldrich^®^, St Louis, USA). Cells were transduced by spinfection for 1 h in the presence of Hexadimethrine bromide (Polybrene) (Sigma Aldrich^®^, St Louis, USA) and media collected and filtered at 48 h and 72 h post-transfection. For human cell lines, constitutive CEP55-knockdown was performed as previously described^1^. The Selection of clones was performed using 400 μg/mL Hygromycin, 50 μg/mL Zeocin, 5 μg/mL Blastocydin (Life TechnologyTM) or 5 μg/mL of Puromycin (Life TechnologyTM). Transient Cep55 silencing was performed by reverse transfection using 10-20 nM of individual small interfering RNAs (siRNAs manufactured by Shanghai Gene Pharma, China) and Lipofectamine RNAiMAX (Life TechnologiesTM) for 48 h.

### Retrovirus and lentivirus packaging and transduction

For the production of retrovirus or lentivirus, Phoenix Amphotropic (retrovirus) or Hek293T cells (lentivirus) were plated at 90 % confluency in a T75 flask and transfected with 5 μg of DNA and 15 μL Polyethylenimine or PEI (Polysciences, Inc., 23966-2, POL) (1:3 ratio) in Optimum media. At 5 h post-transfection, media was changed and the packaging cells incubated for 72 h. At 48 h and 72 h post-transfection, the media was filtered using a 0.45 μm filter onto target cells prior to spinfection (at 1000 X g for 1 h at 25° C) in the presence of hexadimethrine bromide (polybrene; Sigma Aldrich^®^, H9268-5G). Media was removed after spinfection and cells were allowed to recover for the next 48 h before selection with the corresponding antibiotic was carried out to select for transduced cells. Antibiotic selection was sustained until an untransduced control plate of cells had all died.

### Sequences

(5-3’) Cep55_Scr (CCGGCGCTGTTCTAATGACTAGCATCTCGAGATGCTAGTCATT AGAACAGCGTTTTTT); Cep55_sh#2 (CCGGCAGCGAGAGGCCTACGTTAAACTCG AGTTTAACGTAGGCCTCTCGCTGTTTTTG); Cep55_sh#4 (CCGGGAAGATTGAATC AGAAGGTTACTCGAGTAACCTTCTGATTCAATCTTCTTTTTT); SiRNA (Cep55_Scr Sense (5’-3’): CAAUGUUGAUUUGGUGUCUGCA) and anti-sense (5’-3’) : UGAAU AGGAUUGUAAC); SiRNA Cep55_SEQ1 Sense (5’-3’): CCAUCACAGAGCAGCC AUUCCCACT and anti-sense (5’-3’) : AGUGGGAAUGGCUGCUCUGUGAUG GUA)

### Cell and tissue lysate preparation

For preparation of tissue lysate, the organs were diced using a sterile scalpel blade followed by lysis in RIPA buffer (25mM Tris-HCl (pH 7.6), 150mM NaCl, 1% NP40, 1% Sodium deoxycholate and 0.1% SDS) or Urea lysis buffer (8M urea, 1% SDS, 100mM NaCl, 10mM Tris) and the samples were sonicated for 10 seconds on a Branson Sonifier 450 (Branson Ultrasonic Corporation, Danbury, CT, USA). Cell debris was removed by centrifugation at 13,000 RPM at 4°C for 30 minutes. Protein concentration was determined using a Pierce BCA Protein Assay Kit with Bio-Rad Protein Assay Dye Reagent (Thermo-Scientific). 30 or 60 μg of protein was resuspended in 1X laemmli buffer and samples heated to 95°C for 5 minutes prior to electrophoresis.

### Western blot

Polyacrylamide gels were cast as previously described^88^. Prepared protein samples were subjected to electrophoresed at 120 V using the Bio-Rad Mini-PROTEAN^®^ Tetra system in SDS running buffer (25 mM Tris-HCl, 192 mM glycine, 0.1 % SDS (v/v)). Gel transfer was performed using the Invitrogen X-cell SureLock™ transfer system at 80 V for 90 minutes in 1X transfer buffer (50 mM Tris, 40 mM Glycine, 20 % methanol) onto Amersham Hybond nitrocellulose membrane (GE Healthcare, Waukesha, WI, USA), and transfer efficiency assessed by Ponceau S staining (0.1 % (w/v) Ponceau S in 5 % acetic acid). Membranes were blocked in blocking buffer (5 % Skim milk powder (Diploma Brand) in PBS containing 0.5 % Tween-20-PBS-T) for 1 h on a shaker at RT followed by overnight incubation with primary antibodies at 4° C. The membranes were washed (3x PBS-T) and incubated in secondary antibodies for 1 h. Protein detection was performed using Super Signal chemiluminescent ECL-plus (PerkinElmer, Waltham, MA, USA) on a BioRad Gel doc (Bio-Rad ChemiDoc Touch, USA)

### Western blot antibodies

Cep55 (1:1000, In house raised in rabbit against murine Cep55 (amino acids 55-250), Vinculin (1:2000; 13901 Cell Signaling Technology), β-Actin (1:2000, 612656 BD Pharmingen), β-Catenin (1:1000, 9582 Cell Signaling Technology), GSK 3β (1:1000, 9369 Cell Signaling Technology), pGSK 3β(Ser9) (1:1000, 9322 Cell Signaling Technology), Cleaved Caspase-3 (1:500, 9664 Cell Signaling Technology), pAKTs473 (1:1000, 4060 Cell Signaling Technology), AKT (1:1000, 9271 Cell Signaling Technology), p Myc (T58) (1:1000, ab28842 Abcam), Non-phospho (Active) β-Catenin (Ser33/37/Thr41) (1:1000, 8814 Cell Signaling Technology), MYC (Y69) (1:1000, Ab32072Abcam).

